# Extremely acidic proteomes and metabolic flexibility in bacteria and highly diversified archaea thriving in geothermal chaotropic brines

**DOI:** 10.1101/2024.03.10.584303

**Authors:** Ana Gutiérrez-Preciado, Bledina Dede, Brittany Baker, Laura Eme, David Moreira, Purificación López-García

## Abstract

Few described archaeal, and fewer bacterial, lineages thrive at salt-saturating conditions, such as solar saltern crystallizers (salinity above 30%-w/v). They accumulate molar K^+^ cytoplasmic concentrations to maintain osmotic balance (‘salt-in’ strategy), and have proteins adaptively enriched in negatively charged, acidic amino acids. Here, we analyzed metagenomes and metagenome-assembled genomes (MAGs) from geothermally influenced hypersaline ecosystems with increasing chaotropicity in the Danakil Depression. Normalized abundances of universal single-copy genes confirmed that haloarchaea and Nanohaloarchaeota encompass 99% of microbial communities in the near life-limiting conditions of the Western-Canyon Lakes (WCLs). Danakil metagenome- and MAG-inferred proteomes, compared to those of freshwater, seawater and solar saltern ponds up to saturation (6-14-32% salinity), showed that WCL archaea encode the most acidic proteomes ever observed (median protein isoelectric points ≤4.4). We identified previously undescribed Halobacteria families as well as an Aenigmatarchaeota family and a bacterial phylum independently adapted to extreme halophily. Despite phylum-level diversity decreasing with increasing salinity-chaotropicity, and unlike in solar salterns, adapted archaea exceedingly diversified in Danakil ecosystems, challenging the notion of decreasing diversity under extreme conditions. Metabolic flexibility to utilize multiple energy and carbon resources generated by local hydrothermalism along feast-and-famine strategies seemingly shape microbial diversity in these ecosystems near life limits.

---

Extremely halophilic archaea excel in their adaptation to grow in salt-saturating ecosystems, such as solar salterns or athalassohaline hypersaline lakes^1,2^. They include four known lineages recently shown to have independently adapted to halophily^3^: the diverse and long-studied Halobacteria (haloarchaea)^1^, the widespread episymbiotic Nanohaloarchaeota within the DPANN supergroup^4–6^, and the less conspicuous Methanonatronarchaeia^7^ and Halarchaeoplasmatales^8^. Compared to moderate halophiles, which produce compatible solutes to cope with osmotic stress, extremely halophilic archaea accumulate up to 4M K^+^ in their cytoplasm^9,10^. This ‘salt-in’ strategy is concomitant with an excess of acidic amino acids, typically glutamic and aspartic acids, in proteins to preserve their functional structure, such that proteome acidification is a hallmark of extreme halophily^11^. In addition to halophilic archaea, some bacteria and eukaryotes, notably some green algae (*Dunaliella* spp.) and heterotrophic protists, thrive at rather high salt concentrations. However, they use ‘salt-out’ osmoadaptive strategies^10,12^, being absent from saturating environments, such as saltern crystallizer ponds, largely dominated by archaea^13^. The only described exception corresponds to the *Salinibacter* clade, grouping extremely halophilic bacteria mimicking ‘salt-in’ and other archaeal adaptations^14,15^, partly mediated by horizontal gene transfer from haloarchaea^16^.

Most studied hypersaline ecosystems are NaCl-saturated (∼5 M). However, NaCl-dominated brines are thermodynamically moderate^2^ compared to systems of even lower water activity (a_w_) enriched in chaotropic (e.g. Mg, Ca, Li, Fe) salts, which tend to disorganize organic macromolecules^17,18^. Highly chaotropic brines are deleterious and seem devoid of microbial life, such as some hydrothermal brines at and around the Dallol proto-volcano^19,20^. The Dallol area on the Northern Danakil Depression (Afar region, Ethiopia) is situated at the confluence of three major tectonic plates. The local combination of evaporitic and hydrothermal processes^21,22^ produces up-welling thermal fluids enriched in diverse salts and minerals, generating polyextreme brines of contrasting hydrochemistry^23,24^. While some brines across the observed gradients of polyextreme conditions (pH from −1.5 to 6; salinity from ∼30 to >70% w/v; temperature from ∼30°C to 110°C) seem lifeless, others host microbial communities largely dominated by extremely halophilic archaea (up to 99% of community members)^19,25^. In this study, we analyze metagenomes of these hypersaline ecosystems and show that halophilic archaea thriving in the most chaotropic brines permissive for life are remarkably diversified, rely exclusively on heterotrophic processes and push known adaptations to unprecedented limits.

## Results and Discussion

### Proteome-wide adaptation of archaea and bacteria in increasingly chaotropic brines

We sequenced, assembled and annotated metagenomes from microbial communities thriving in geothermally influenced hypersaline systems in the north Danakil salt desert (Fig.1a; Supplementary Table 1). These included two locations from Lake Karum or Assale sampled in different years (Ass, 9Ass), one cave reservoir at the Dallol proto-volcano salt canyons (9Gt) and two of the Western-Canyon Lakes (WCL2, WCL3). WCL3 displayed the highest salinity and the lowest water activity (a_w_) and pH along the sampled gradient^19,25^ (Fig.1a). The WCLs had the highest joint Ca^2+^+Mg^2+^ concentrations, followed by Lake Assale samples (Supplementary Table 2), making these systems highly chaotropic^19,25^. The microbial community composition inferred from the normalized abundance of selected universal single-copy genes (USCGs; Supplementary Fig.1) in metagenomes was dominated by archaea, overwhelmingly so in the WCLs (Fig.1b), consistent with previous 16S/18S rRNA gene metabarcoding studies^19,25^. Members of the class Halobacteria and the phylum Nanohaloarchaeota were by far the most abundant archaea. They encompassed widely diverse genera that showed different distribution patterns among samples, particularly marked between Assale and WCL samples (Extended Data Fig.1). Bacteria were relatively diverse, with photosynthetic cyanobacteria detected only in Lake Assale, where Salinibacteraceae were also relatively abundant (Fig.1b). Eukaryotic sequences were negligible to undetectable, notably in WCLs, also confirming previous observations^19,25^.

**Fig.1.**
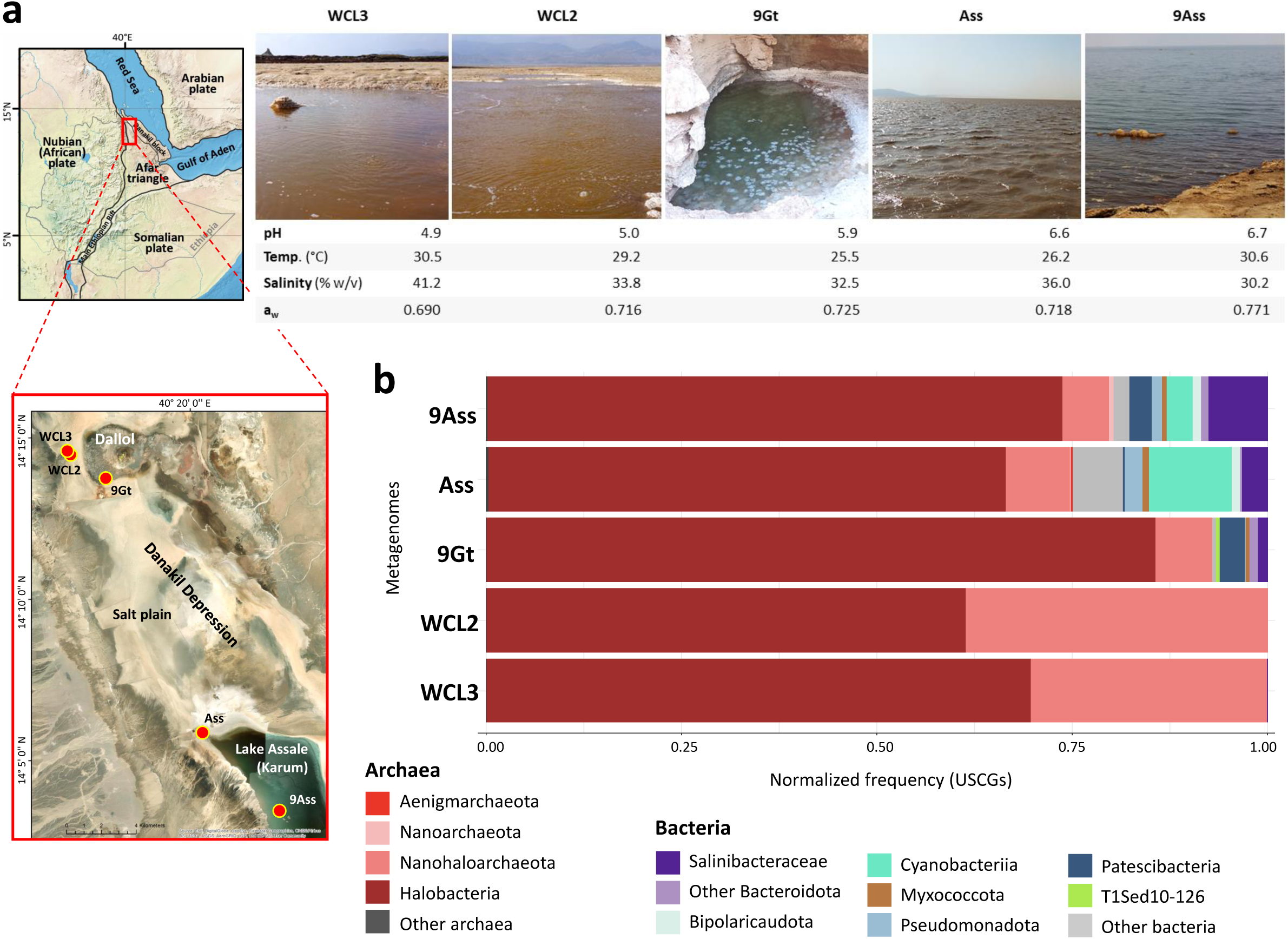
Microbial community composition inferred from metagenomes of polyextreme chaotropic ecosystems in the north Danakil Depression. **a,** Sampling sites around the Dallol proto-volcano and Lake Assale or Karum in the north Danakil Depression, Ethiopia; some major abiotic parameters for each system are shown (a_w_, water activity). **b**, Global microbial community composition at high-rank taxonomic level of sampled polyextreme ecosystems inferred from the normalized frequency of universal single-copy genes (USCGs; selection of ribosomal proteins expressed in RPKM). Note that DAL-WCL2 and WCL3 are composed of 99% archaea (Halobacteriota and Nanohaloarchaeota; classification according to GTDB r214).

We investigated known proteome-wide adaptations to extreme halophily at metagenome level (Fig.2). First, we calculated the isoelectric point (pI) of proteins encoded by the Danakil metagenomes in comparison to those from ecosystems of increasing salinity: a freshwater lake in France, Mediterranean seawater samples and solar saltern ponds containing 6%, 14% and 32% salt (w/v), the latter relatively enriched in Mg^2+^; Supplementary Table 2). As expected^26^, samples up to 14% salt displayed a bimodal distribution, with some acidic and some basic proteins, while the 32%-salt saltern pond displayed a unimodal distribution with a marked peak at acidic pH (median pI, 4.55; Supplementary Table 2). The Danakil inferred proteomes also displayed a unimodal distribution, with Lake Assale exhibiting similar values to those of the 32%-salt pond. However, the WCLs exhibited a more pronounced peak shifted to even more acidic values (median pI∼4.4) (Fig.2a). This reflected a strong amino acid bias, with aspartic and glutamic acids (D+E) being enriched and isoleucine and lysine (I+K) depleted in Danakil metagenomes, a well-known adaptation to extreme halophily^11^. The DE/IK ratio was much higher in the WCL metagenomes, concurrent with the higher [Ca^2+^] and [Mg^2+^] of these brines (Fig.2b-c). We also observed a marked preference, among positively charged amino acids, for arginine versus lysine in proteomes from environments displaying increasing salt concentrations, with the R/K ratio being also highest in the WCLs (Fig.2c; Extended Data Fig.2). This is partly due to the usually high GC content of haloarchaeal genomes: arginine codons are GC-rich whereas lysine codons are AT-rich^27^. This trend extends to all GC-enriched and depleted codons, corresponding to GARP and FIMNKY amino acid groups, respectively^3^. In addition, arginine is further favored over lysine due to its higher coil forming propensity^27^ and the ability to bind more water molecules along its lateral chain, which helps maintain a hydrated protein state^28,29^.

**Fig.2.**
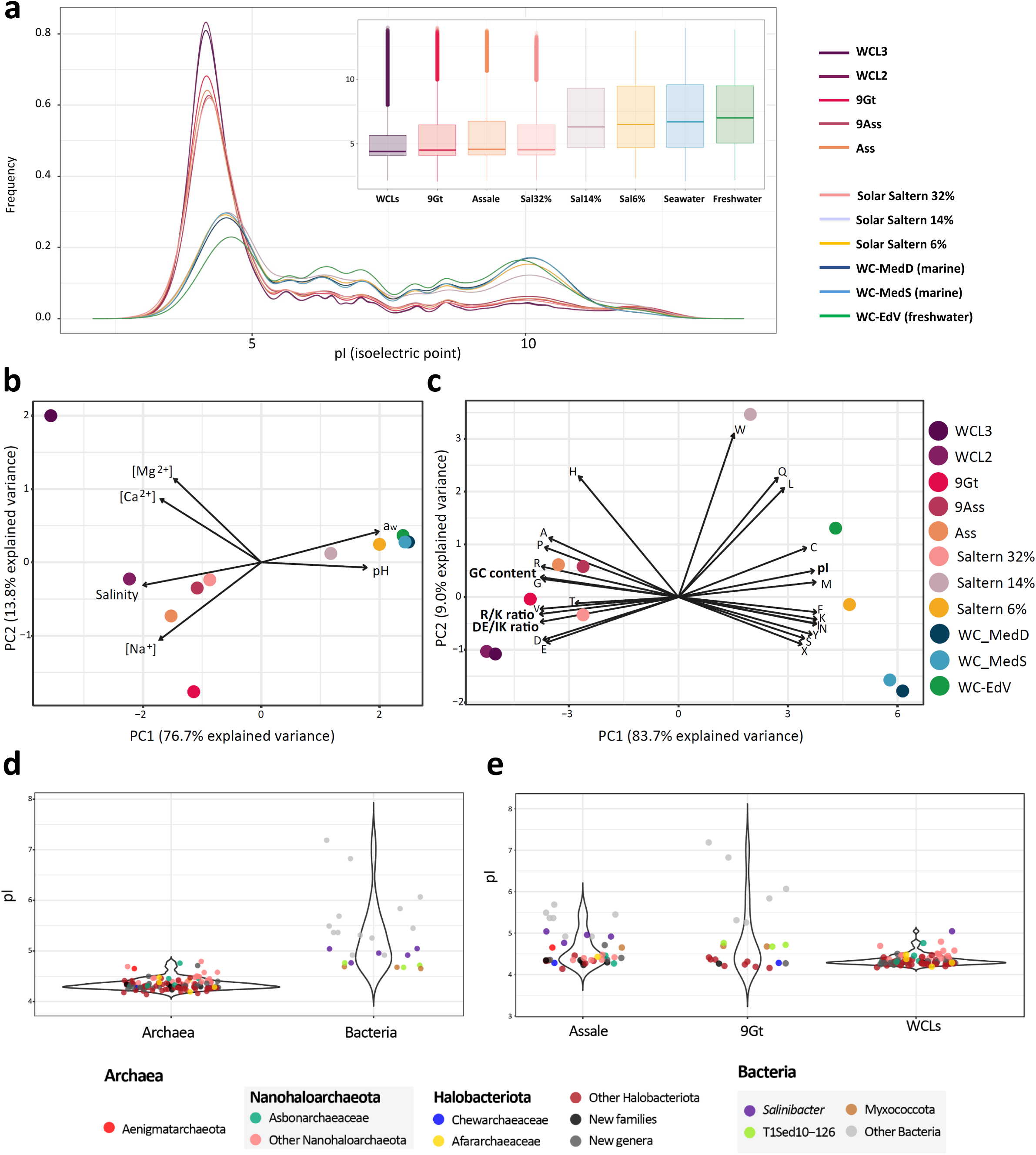
Isoelectric point and amino acid compositional biases of inferred proteomes for microorganisms thriving in increasingly chaotropic ecosystems from the north Danakil Depression. **a,** Distribution of isoelectric point (pI) values inferred for proteins encoded by the analyzed Danakil metagenomes in comparison with representative metagenomes from freshwater, seawater and solar saltern brines of increasing salt concentration (6-14-32%). Values for reference ecosystems were inferred from two replicate metagenomes each. The inlet shows a barplot displaying pI values per ecosystem type. **b**, Principal component analysis (PCA) of the Danakil polyextreme systems and ecosystems sampled along a salinity and chaotropicity gradient as a function of major abiotic determinants; chaotropicity is associated with high Mg^2+^ and Ca^2+^ concentrations. **c**, PCA of the GC content and inferred pI median values, amino acid composition and DE/IK and R/K ratios from the analyzed metagenomes. **d**, jitter-violin plot showing protein pI values inferred from individual metagenome-assembled genomes (MAGs) affiliating to the domains Archaea and Bacteria. **e**, jitter-violin plot showing proteome pI values inferred from MAGs in Danakil brine systems.

These adaptations were also clearly visible at the level of individual MAGs and their affiliated taxa. As expected, archaeal MAGs displayed the lowest pI values as compared to the few bacterial MAGs retrieved (Fig.2d). Interestingly, MAGs affiliating to marine uncultured Myxococcota (genus CAJXPB0, Bradymonadaceae) and the phylum T1Sed10-126 exhibited pI values comparable to the Salinibacteraceae (Fig.2d; Supplementary Table 3), suggesting the occurrence of similar adaptations to hypersaline environments in bacteria other than the Salinibacteraceae. Average pI values of MAGs were clearly lower in the WCLs as compared to Lake Assale and 9Gt samples (Fig.2e) and their respective pI distributions were also more shifted towards unimodal low pI distributions (Extended Data Fig.3a). This reflects the stronger selection pressure exerted by the near life-limiting WCL hypersaline chaotropic conditions. Despite the acidic pI values, amino acid biases were slightly different depending on the taxon. Thus, Halobacteria MAGs displayed the highest DE/IK and R/K ratios, except for the recently described deep-branching Afararchaeaceae family^3^, which showed a less biased amino acid content, more comparable to that of Nanohaloarchaeota MAGs (Extended Data Figs.2b and 3c-d).

### Expanded diversity of extremely halophilic lineages

It is generally believed that microbial diversity decreases as the physicochemical conditions approach those that are limiting for life^30,31^. This seems valid at least at high-rank taxon level. For instance, only members of the archaeal domain, and from a limited number of phyla, optimally thrive at temperatures higher than 95°C^32–34^. Likewise, as illustrated by the Danakil systems studied here, only members from a small number of archaeal phyla or classes thrive in salt-saturated brines^1,4,7,8^, with extremely halophilic bacteria being anecdotal in hypersaline chaotropic systems, such as the WCLs (Figs.1-2). To investigate whether this restricted diversity is also observable at finer taxonomic scale, we generated operational taxonomic units (OTUs) based on clusters of USCGs and determined their normalized relative abundance across metagenomes from increasingly salty ecosystems. We observed a decrease in global alpha diversity and Shannon and Simpson diversity indexes in 14% and, most especially, 32% salinity ponds compared to freshwater, marine and 6%-salt solar saltern ponds (Fig.3; Supplementary Fig.2a). The diversity indexes decreased in the 32%-salt pond relative to that of 14%-salt even for Halobacteria, as a consequence of the overdominance of *Haloquadratum* spp. in the 32%-salt pond analyzed metagenomes^26,35^ (Supplementary Fig.2b). In contrast, global alpha diversity and diversity indexes were much higher in the Danakil brines, and comparable to those of non-extreme ecosystems, the key distinction being that, in the Danakil hypersaline chaotropic systems, the Halobacteria diversity virtually represented the total ecosystem diversity. A similar trend, albeit to a lesser extent, was observed for the Nanohaloarchaeota diversity parameters (Fig.3a-b). This implies that, after they evolved the necessary adaptations to thrive in hypersaline environments, haloarchaea radiated greatly, secondarily adapting to a variety of additional environmental constraints and ecological niches.

**Fig.3.**
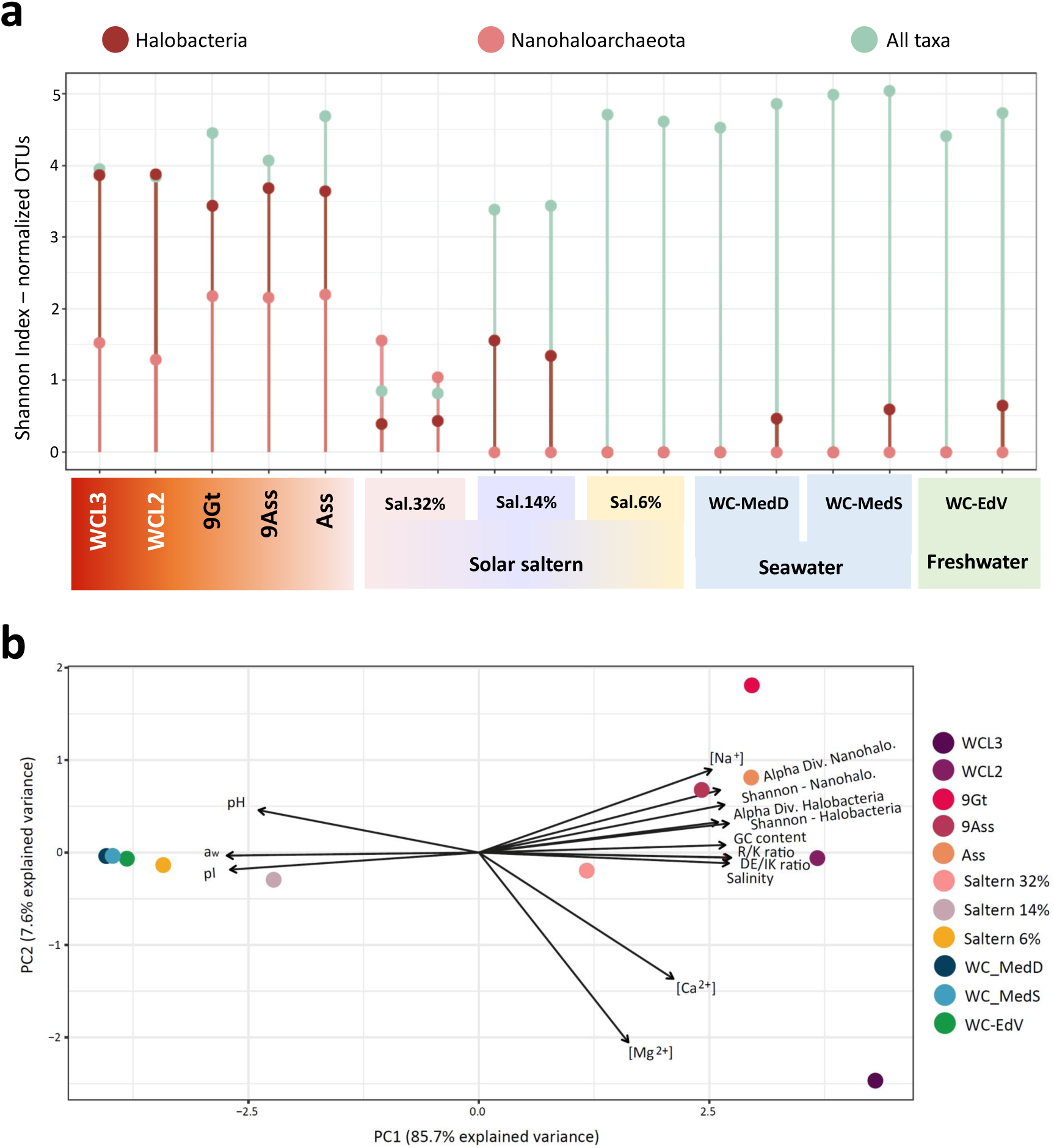
Diversity indexes for all species and dominant archaeal taxa in Danakil geothermally influenced chaotropic brines compared to other ecosystems of increasing salinity. **a,** Shannon index calculated from normalized USCGs from individual metagenomes grouped at species-level operational taxonomic units (OTUs; see Methods) considering OTUs for all taxa, Halobacteria and Nanohaloarchaeota. **b**, PCA of various compositional biases, abiotic factors and alpha-diversity and Shannon diversity indexes for dominant archaeal taxa in Danakil polyextreme brines as compared to reference ecosystems along a salinity gradient.

This wide multiplicity of extremely halophilic archaea in the Danakil ecosystems was partially captured by the phylogenetic diversity of assembled MAGs. We could assemble 483 MAGs, from which 155 were more than 40% complete and less than 5% redundant. The relatively low recovery of complete MAGs was partly due to the high genomic diversity. We classified these MAGs based on both, the Genome Taxonomy Database (GTDB^36^) taxonomy classifier^37^, as well as Maximum Likelihood (ML) phylogenomic analyses using a curated dataset of 127 non-ribosomal protein markers for archaea that yields robust phylogenies^3^ (Fig.4). Out of the 155 most complete MAGs, 92 affiliated to the Halobacteria, 38 to the DPANN supergroup and 25 to bacteria (Supplementary Table 3). Based on a phylogeny that included a representative of each described genus of the class Halobacteria combining this information, we not only identified diverse members in the families Halobacteriaceae, Haloferacaceae and Haloarculaceae, with several potential new genera, but also up to four additional Halobacteria families (Fig.4a; Supplementary Table 3). Two of them were recently described from these Danakil ecosystems, the Afararchaeaceae and the Chewarchaeaceae^3^. In our phylogenomic tree, *Halorutilus salinus*, the type species of a recently described new family^38^, branched within the Afararchaeaceae, which therefore becomes a junior synonym for the family Halorutilaceae (Fig.4a). We additionally identified two other family-level clades that we name Karumarchaeaceae and Abyssiniarchaeaceae, having as representative genomes, respectively, the MAGs DAL-Ass_21_92C4R, described as *Karumarchaeum halophilus*, and DAL-8mg-83m_91C4R, as *Abyssiniarchaeum dallolvicinus* (estimated genome sizes of 2.6 and 2.4 Mbp, respectively; see formal description below). Curiously, we did not detect any member of other Halobacteria families, several of which include alkaliphilic members. This suggests that the conditions of these highly chaotropic ecosystems are not conducive to their development, perhaps due to the slightly acidic conditions (pH∼5-6.7). We also identified many Nanohaloarchaeota members, including potential new genera, within the Nanosalinaceae and the recently described family Asbonarchaeaceae from these Danakil ecosystems^3^ (Fig.4b).

**Fig.4.**
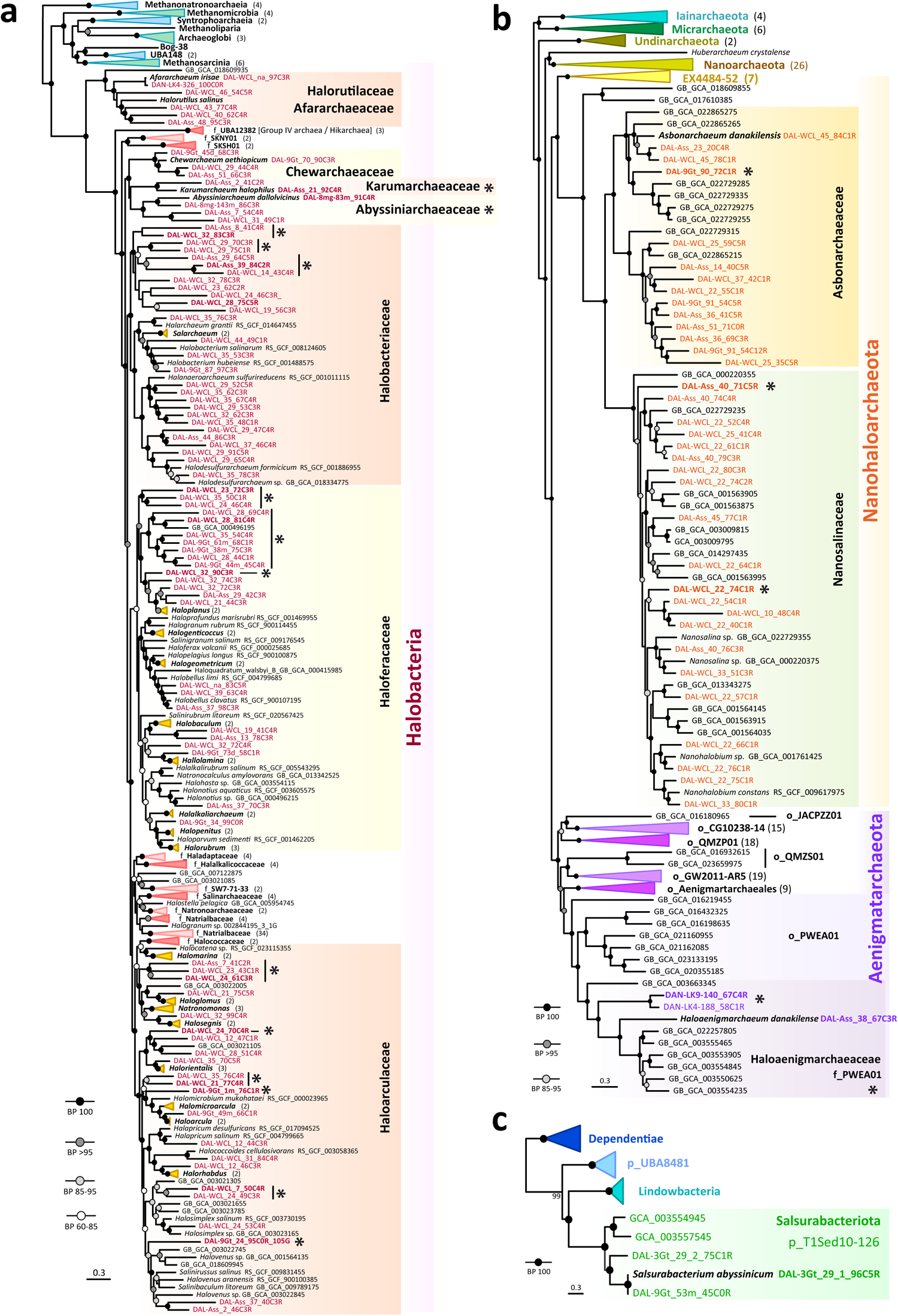
Phylogenomic trees of newly identified archaeal and bacterial MAGs in hypersaline chaotropic north Danakil ecosystems. **a,** Phylogenomic tree of archaeal MAGs affiliating to the class Halobacteria (Halobacteriota). **b**, Phylogenomic tree of MAGs affiliating to the Nanohaloarchaeota and Aenigmatarchaeota (DPANN supergroup). **c**, Phylogenomic analysis of the bacterial phylum *Candidatus* Salsurabacteriota (p_T1Sed10-126), grouping seemingly extreme halophilic members. Bootstrap value ranges are indicated at nodes. Names of MAGs assembled from our Danakil metagenomes are highlighted in color; reference sequences are indicated in black. Some taxa were collapsed to facilitate visualization of trees (detailed trees are shown in Supplementary Figs. 4-6); the number of total representatives included in collapsed taxa is given in brackets. Asterisks indicate genera and families newly identified in this study.

Interestingly, the MAG DAL-Ass_38_67C3R branched within the Aenigmarchaeota family f_PWEA01 (Fig.4b) and exhibited the characteristic pI unimodal distribution of extreme halophiles (Extended Data Fig.4a). Two other Danakil MAGs branching more deeply within the family, also showed similar pI distributions, albeit slightly less pronounced. GTDB genomes belonging to f_PWEA01, but not to other families of the same order (o_PWEA01), also display unimodal acidic pI distributions (Extended Data Fig.4a). These genomes were retrieved from hypersaline soda lake sediments (Kulunda Steppe, Altai, Russia)^39^. Collectively, this strongly suggest that the f_PWEA01 represents a lineage of archaea independently adapted to hypersaline conditions, which we propose to rename as Haloaenigmatarchaeaceae (type species *Haloaenigmatarchaeum danakilense*, represented by the MAG DAL-Ass_38_67C3R, of ∼1.5 Mbp). Likewise, three Danakil MAGs branched within the bacterial GTDB phylum p_T1Sed10-126 (Fig.4c), originally defined by two GTDB genomes retrieved from the same hypersaline lake sediment in Siberia^39^. This genomic clade, as other newly detected lineages mentioned above, was cohesive, as suggested by average nucleotide identity values^40^ (Supplementary Fig.3). These five genomes also display biased pI distributions, with median pI values lower than 5, similar to the Salinibacteraceae (Extended Data Fig. 4b-c). This suggests that the bacterial phylum p_T1Sed10-126, which we have renamed Salsurabacteriota (type species *Salsurabacterium abyssinicum*, represented by the MAG DAL-3Gt_29_1_96C5R) independently adapted to extreme halophily.

Why is the diversity of extremely halophilic archaea so high in the increasingly chaotropic brines of the Northern Danakil compared to the apparently less harsh conditions of solar saltern salt-saturating ponds? We hypothesize that it is associated with richer resource availability due to the local geochemical settings and the occurrence of diverse metabolic capacities to exploit those resources.

### Metabolic flexibility and feast-and-famine strategies

The hydrothermal activity affecting the Danakil ecosystems not only provides an input of reduced gases, which can serve as potential electron donors, but are also enriched in organics derived from the interaction of mantle fluids and Proterozoic sediments below the salt-crust^21,41^. Chemical analyses showed the presence of diverse organics in the Dallol area brines, including the WCLs^19,25^. To determine whether these resources could sustain the observed microbial communities, we characterized their metabolic potential using complementary approaches. We first carried out general metabolic inferences based on diagnostic functional genes and KEGG modules detected in the reconstructed MAGs and several genomes from their closest neighbor and outgroup taxa. This showed that Halobacteria members were heterotrophs possessing classical core biosynthetic functions for amino acids, nucleotides and lipid components, alongside capabilities for aerobic and anaerobic respiration (Extended Data Fig.5). Different MAGs/taxa encoded genes to utilize different organics: diverse, including branched, amino acids, hydrocarbons, fatty acids or aromatic compounds. Thus, in addition to core functions such as the Krebs cycle, glycolysis, gluconeogenesis and biosynthesis of amino acids and nucleotides, Karumarchaeaceae and Abyssiniaceae shared aerobic and anaerobic (nitrite and CO) respiration, fermentation, and chlorite and iron/manganese reductive capabilities. However, whereas *Abyssiniarchaeum* seemed to rely mostly on amino acid and fatty acid degradation, *Karumarchaeum* likely uses diverse complex hydrocarbons, encoding endohemicellulases, and amylotytic and cellulose-degrading enzymes (Extended Data Fig.5). By contrast, the metabolic potential of Nanohaloarchaeota MAGs was considerably reduced, having missing or incomplete essential biosynthetic pathways (Extended Data Fig.6), which suggests that they depend on their haloarchaeal hosts for survival^5,6^. They seem able to ferment and, like other Nanohaloarchaeota, have an ATP-synthase despite the absence of an identifiable electron-transport chain. Intriguingly, several MAGs possessed amylolytic-type enzymes (Extended Data Fig.6a). This opens the possibility that some nanohaloarchaea’s parasitic relationships skirt on the edge of mutualism along the symbiotic spectrum. They could provide metabolic complementation to their hosts for the degradation of specific hydrocarbons in exchange for numerous essential compounds, as has been already observed in some haloarchaea-nanohaloarchaea consortia^42,43^. This peculiar mutualism could explain the apparently stable prevalence of Nanohaloarchaeota in these ecosystems, notably the WCLs (close to 40%, Fig.1), without leading to parasitic overload and population collapse. Members of the bacterial phylum Salsurabacteriota, similar to classical haloarchaea, encoded core metabolic pathways, being likely able to respire aerobically and anaerobically and to use amino acids and fatty acids (Extended Data Fig.7).

Second, to determine the primary electron acceptors preferentially used by the Danakil microbial communities, we investigated and manually verified the presence of diagnostic genes involved in energy-transducing redox reactions in both, metagenomes and MAGs (Fig.5). We focused on redox processes leading to the reduction of oxygen, nitrogen and sulfate. In dynamic microaerophilic conditions, like those encountered in these geothermally influenced Danakil lakes, especially in the actively degassing WCLs, microorganisms adapt by either utilizing cytochromes with a high affinity for nanomolar O_2_ levels^44^ or alternative compounds such as organic molecules, nitrogen derivatives, or sulfur species. Oxygen respiration genes, including low and high-affinity cytochromes, were consistently abundant in all metagenomes, indicating adaptation to varying O_2_ concentrations^44^ (Fig.5). Cytochromes were present across all MAGs recovered in this study, except for those of Nanohaloarchaeota. Nitrate respiration is adaptive in hypersaline ecosystems, especially at relatively high temperatures, due to the low oxygen solubility under these conditions^11^. Indeed, genes encoding nitrate reductases, which catalyze the reduction of nitrate to nitrite, were far more prevalent than oxygen respiration genes. They were widespread across all metagenomic samples, being encoded in up to 20% of genomes in Lake Assale. They occurred in MAGs from virtually all the detected Halobacteria families, and in *Salsurabacterium* (Fig.5). Other genes associated with the nitrogen cycle were also highly prevalent. The isotopic signature of nitrogen compounds in the Dallol area volatiles suggests an important input of mantle-derived N sources^41^. Nitrate can oxidize organic matter^45^, methane^46,47^, sulfur compounds^48^ or iron^49^, in addition to being a source of nitrogen^50^. The conversion of nitrite to ammonium is also utilized for both dissimilatory and assimilatory processes^50^. Ferredoxin-dependent assimilatory nitrite reductase was relatively abundant in all Danakil metagenomes and occurred in *Karumarchaeum*, Haloferacaceae and Haloarculaceae MAGs (Fig.5; Supplementary Table 4), further supporting that nitrate and nitrite are primary nitrogen sources. Additionally, genes responsible for denitrification, which facilitate the sequential conversion of nitrite to nitric oxide, nitrous oxide, and ultimately nitrogen gas, were also prevalent. Genes encoding copper-dependent nitrite reductases, converting nitrite to nitric oxide, nitric oxide and nitrous oxide reductases were present in several Halobacteria MAGs and *Salsurabacterium* (Fig.5). Assimilatory nitrite reduction and denitrification are well known in haloarchaea^51^. Genes related to sulfur and fumarate reduction were as abundant as aerobic respiration-related cytochromes, again indicative of low O_2_ levels. Fumarate reductase, used in reducing fumarate to succinate was present in almost all MAGs, while sulfate adenylytransferase (Sat) was present in *Karumarchaeum*, Halobacteriaceae, *Salsurabacterium* and even Nanohaloarchaeota, but was missing in Haloferacaceae MAGs. Given the geothermal settings, we investigated genes involved in chemolithotrophy, i.e. energy-generating redox reactions involving the oxidation of inorganic compounds, including sulfur reduced species, hydrogen, and carbon monoxide. Among these, CO oxidation appeared the most prominent in the Dallol area lakes, with CO-dehydrogenase encoded in 3-7% of genomes from these ecosystems (Fig.5). However, the genes for CO and hydrogen oxidation were not identified in any of the retrieved MAGs, suggesting that are distributed in diverse, less dominant microorganisms. Sulfur oxidation genes (sulfide dehydrogenase and sulfide:quinone oxidoreductase), were present in *Abyssiniarchaeum*, *Karumarchaeum*, Halobacteriaceae, Haloferacaceae and Haloarculaceae, reinforcing the idea that geothermal activity significantly influences the microbial communities in these polyextreme lakes.

**Fig.5.**
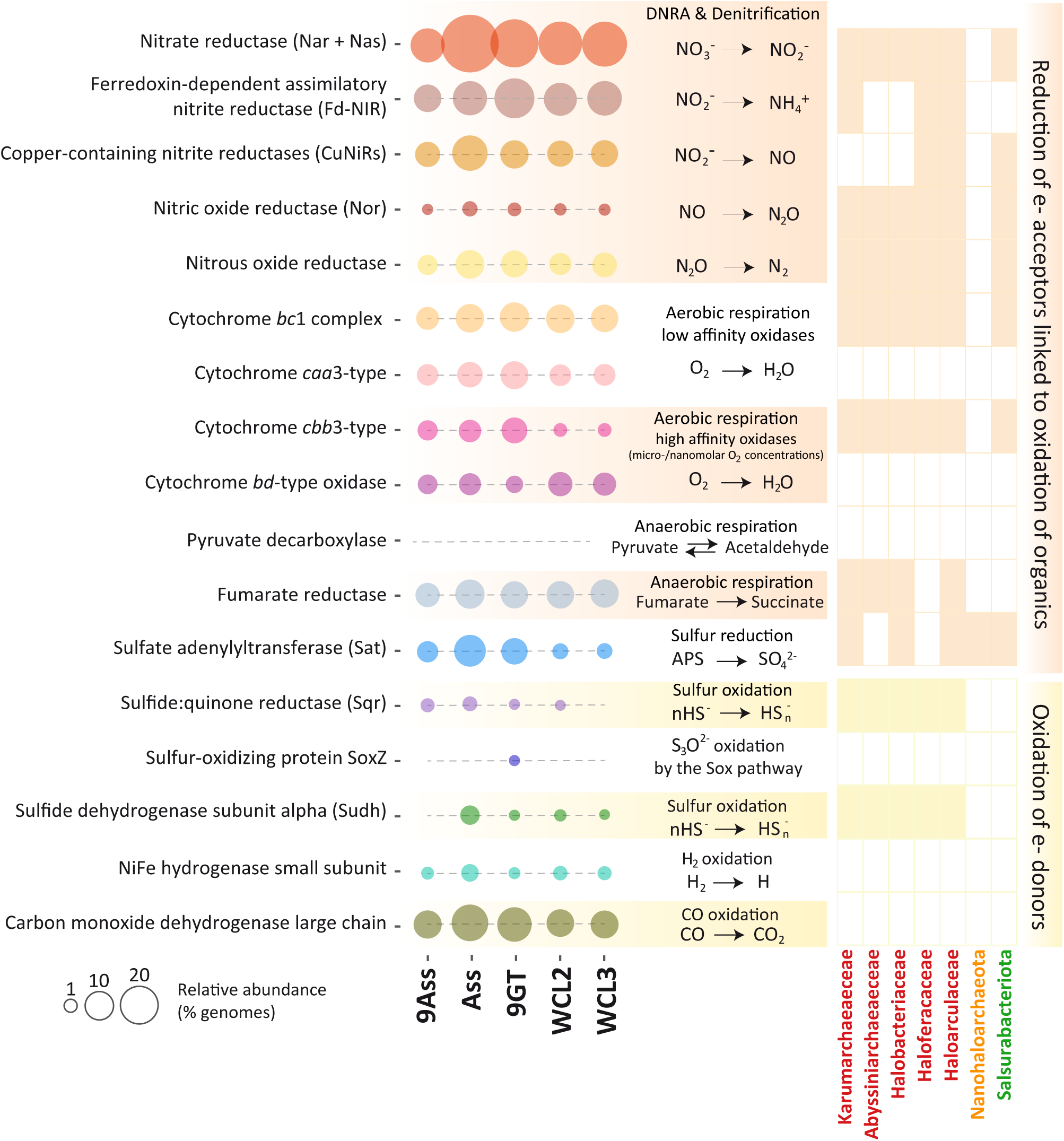
Major processes involved in energy metabolism in microbial communities from Danakil polyextreme brine ecosystems. The relative abundance of genes in the different metagenomes, shown on the left, is expressed in terms of the inferred proportion of genomes harboring them. The presence absence of the corresponding genes in MAGs of the selected taxa is shown on the right. DNRA, Dissimilatory Nitrate Reduction to Ammonium. APS, 5′-adenylsulfate.

These Danakil communities seem to largely rely on heterotrophic processes. They apparently lack chemosynthetic carbon fixation and the WCLs also lack photosynthetic members^25^ (Fig.1). Since, in addition to amino acid and fatty acid degradation, hydrocarbon utilization appeared important (Extended Data Fig.5), we searched for glycoside hydrogenase (GH) and alkane degradation genes. In Lake Assale samples (Ass, 9Ass) and 9Gt, the most prevalent GH genes participate in the degradation of starch/glycogen and mixed polysaccharides (GH15, GH29, and GH3^52,53^; Extended Data Fig.8). In addition, the consistent presence of haloalkane dehalogenases and alkane monooxygenase across metagenomes suggests an input of haloalkanes and/or short-chain alkanes, consistent with the analysis of volatiles in some Dallol area lakes^41^. However, flavin-binding monooxygenase genes, known to be involved in the breakdown of long-chain molecules^54^ were rare across samples except for Lake Assale, suggesting limited capability to decompose long-chain carbon molecules (+32C) in these ecosystems. The diversity and abundance of GH genes increased with salinity-chaotropicity, especially in the WCLs (Extended Data Fig.8), suggesting that these extremely halophilic archaea and possibly also bacteria^55^ degrade a broad range of hydrocarbons^56^, mirroring observations in Ethiopian soda lakes^57^. In addition, Danakil MAGs contained genes for carbon storage, such as polyhydroxybutyrate biosynthesis genes (phbC), a common trait in many haloarchaea^58^.

The ability, inferred in several MAGs, to degrade a wealth of polysaccharides, store carbon and reduce oxygen, nitrate, nitrous oxide, nitric oxide, sulfate and fumarate, points towards a “feast-or-famine” metabolic strategy. This is an adaptive response to fluctuating environmental conditions, whereby microorganisms must rapidly exploit available resources and be able to survive periods of scarcity. Feast-or-famine strategies are observed in energy-depleted environments, which occasionally (periodically or spatially) receive inputs of nutrients, such as the deep sea^59^. During “famine” periods, microorganisms derive energy from stored carbon storage or the degradation of recalcitrant organic matter. In the Danakil hypersaline ponds, “famine” periods could correspond to dominant evaporative phases leading to full desiccation or trapping in halite brine inclusions^60^. “Feast” phases could be triggered by the geothermal activity linked to meteoric waters infiltrating from the high Ethiopian plateau towards the depression and the concomitant generation of upwelling fluids^21,22,24^, providing hydration and nutrients. The high variety of exploitable metabolic resources and abilities may partly explain the diversification of extremely halophilic archaea in these chaotropic Danakil ecosystems. Since microbial diversity depends not only on the accessibility to various resources but also in resource partitioning through trophic interactions^61,62^, the combination of multiple resources linked to the local geothermal context amplified by trophic network interactions likely drive the adaptation to a myriad ecological niches and, consequently, the observed diversity.

### Conclusions

We analyzed metagenomes of several hypersaline and increasingly chaotropic ecosystems influenced by geothermal activity in the Dallol area, Northern Danakil Depression, Ethiopia. Some of these ecosystems, notably the WCLs, were the most polyextreme environments sampled in the area harboring microbial life^19,20,25^. We showed that these ecosystems were overwhelmingly dominated by extremely halophilic Halobacteria and Nanohaloarchaeota but, contrary to expectations suggesting that low diversity associates with increasingly extreme conditions, we observed an unprecedented diversity of archaea. They adapt to the challenging osmotic conditions via a ‘salt-in’ strategy and have record-acidic proteomes when compared with archaea thriving in aquatic environments of increasing salinity (freshwater to 32%-salinity solar saltern ponds). We uncovered several family-level and genus-level clades of Halobacteria and Nanohaloarchaeota previously undescribed, as well as a new family of Aenigmarchaeota (Haloaenigmatarchaeaceae), which likely represents a fifth independent convergent adaptation to extreme halophily in the archaeal domain^3^. Additionally, we identified a new phylum of extreme halophilic bacteria, the Salsurabacteriota (p_T1Sed10-126), displaying proteomes as enriched in acidic amino acids as the halophilic Salinibacteraceae. Metagenome and MAG analyses allow to infer that these microbial communities rely on a wide variety of carbon sources and electron donors and acceptors. In particular, the WCLs likely constitute fully heterotrophy-based ecosystems depending on organics largely mobilized by upwelling hydrothermal fluids interacting with Proterozoic marine sediments below the Danakil desert salt crust^21,22,24^. Alkane and haloalkane degradation seem particularly important resources for these communities. Collectively, the vast array of carbon sources and redox reactions combined with resource partitioning through trophic networks can explain the unprecedented diversity observed in these microbial communities thriving close to life-limiting boundaries.

### Taxonomic descriptions

All new taxa have been described under the SeqCode^63^ as follows:

***Karumarchaeum* gen. nov. Etymology.** *archaeum* (N.L. neut. n.): an archaeon; *Karumarchaeum* (N.L. neut. n.): an archaeon from Lake Karum, Afar region, Ethiopia. Type species, *Karumarchaeum halophilus*.

***Karumarchaeum halophilus* sp. nov. Etymology**. *halophilus* (N.L. masc. adj.): salt-loving.

**Diagnosis**. This archaeon lives in in suboxic hypersaline waters influenced by hydrothermal activity. It encodes for aerobic and anaerobic respiration, including denitrification. It is able to use amino acids and likely relies on halogenated compounds as well as cellulose and other complex hydrocarbon polymers for organo- and/or chemo-heterotrophic growth. Its genome is around 2.6 Mbp (GC content: 62%). It is known from environmental sequencing only. The designated type MAG is DAL-Ass_21_92C4R.

**Karumarchaeaceae fam. nov. Etymology**. *Karumarchaeum* (N.L. neut. n.): a genus name; - aceae, ending to denote a family; Karumarchaeaceae (N.L. fem. pl. n.): the *Karumarchaeum* family.

***Abyssiniarchaeum* gen. nov. Etymology.** *archaeum* (N.L. neut. n.): an archaeon; *Abyssiniarchaeum* (N.L. neut. n.): an archaeon from Abyssinia, former name of the Ethiopian Empire. Type species, *Abyssiniarchaeum dallolvicinus*.

***Abyssiniarchaeum dallovicinus* sp. nov. Etymology**. *dallolvicinus* (N.L. masc. adj.): neighboring the Dallol proto-volcano in the north Danakil Depression.

**Diagnosis**. This archaeon lives in suboxic hypersaline environments influenced by hydrothermal activity. It encodes for aerobic and anaerobic respiration. It can use amino acids and likely relies on halogenated compounds, fatty acids and some hydrocarbons for organo- and/or chemo-heterotrophic growth. Its genome is around 2.4 Mbp (GC content: 67%). It is known from environmental sequencing only. The designated type MAG is DAL-8mg-83m_91C4R.

**Abyssiniarchaeaceae fam. nov. Etymology**. *Abyssiniarchaeum* (N.L. neut. n.): a genus name; -aceae, ending to denote a family; Abyssiniarchaeaceae (N.L. fem. pl. n.): the *Abyssiniarchaeum* family.

***Haloaenigmatarchaeum* gen. nov. Etymology.** *archaeum* (N.L. neut. n.): an archaeon; *Haloaenigmatarchaeum* (N.L. neut. n.): a salt-loving archaeon of the phylum Aenigmatarchaeota). Type species: *Haloaenigmatarchaeum danakilense*.

***Haloaenigmatarchaeum danakilense* sp. nov. Etymology**. *danakilense* (N.L. neut. adj.): pertaining to the Danakil Depression.

**Diagnosis**. This archaeon lives in hypersaline systems of the Danakil Depression. It has a reduced genome of around 1.5 Mbp (GC content: 48%) and lacks most essential biosynthetic pathways, most likely growing as a symbiont of an unknown host. It is known from environmental sequencing only. The designated type MAG is DAL-Ass_38_67C3R.

**Haloaenigmatarchaeaceae fam. nov. Etymology**. *Haloaenigmatarchaeum* (N.L. neut. n.): a genus name; -aceae, ending to denote a family; Haloaenigmatarchaeaceae (N.L. fem. pl. n.): the *Haloaenigmatarchaeum* family.

***Salsurabacterium* gen. nov. Etymology**. *bacterium* (N.L. neut. n.): a bacterium; *Salsurabacterium* (N.L. neut. n.): a bacterium thriving in brine. Type species: *Salsurabacterium abyssinicum*.

***Salsurabacterium abyssinicum* sp. nov. Etymology**. *abyssinicum* (N.L. neut. adj.): pertaining to Abyssinia, former name of the Ethiopian Empire.

**Diagnosis**. This bacterium thrives in brines of the Danakil Depression, Ethiopia. It is likely capable of aerobic and anaerobic respiration and organoheterotrophic growth. Its genome has 3.8 Mbp (GC content: 51%). It is known from environmental sequencing only. The designated type MAG is DAL-3Gt_29_1_96C5R.

**Salsurabacteriota phyl. nov. Etymology**. *Salsurabacterium* (N.L. neut. n.): a genus name; - ota, ending to denote a phylum; Salsurabacteriota (N.L. neut. n.): the *Salsurabacterium* phylum).

## Materials & Methods

### Danakil samples, DNA purification and metagenome sequencing

Northern Danakil brine samples used for metagenome sequencing were collected from Lake Karum or Assale in 2016 (Ass) and 2019 (9Ass) and from an underground cave reservoir in the Dallol canyons (La Grotte, 9Gt) and the Western Canyon Lakes (WCL2, WCL3) in 2019. The specific description of these sites, their hydrochemistry and the associated microbial community composition based on 16S rRNA gene amplicon metabarcoding studies has been published^19,25^. The main physicochemical parameters and cation concentrations are highlighted in Fig.1 and Supplementary Table 2. Brine samples (5-25 l) were sequentially filtered through 30-µm and 0.22-µm pore-diameter Nucleopore filters (Whatman, Maidstone, UK) and the filters retaining the 0.2–30 µm cell fraction were fixed with absolute ethanol (>80% final concentration) in 2-ml cryotubes and stored at −20°C until use. After ethanol elimination and biomass rehydration, DNA was purified using the Power Soil DNA Isolation Kit (MoBio, Carlsbad, CA, USA) under a UV-irradiated Erlab CaptairBio DNA/RNA PCR Workstation. DNA was resuspended in 10 mM Tris-HCl, pH 8.0 and stored at −20°C. Total DNA was sequenced using HiSeq Illumina paired-end (2×125 bp) by Eurofins Genomics (Ebersberg, Germany). Metagenome statistics and GenBank accession numbers are provided in Table S1.

### Sequence analysis, functional annotation, and metagenome-inferred microbial community composition

Raw Illumina reads were quality verified with FastQC v0.11.8 and cleaned with Trimmomatic^64^ v0.39, adjusting the parameters as needed (usually LEADING:3 TRAILING:3 MAXINFO:30:0.8 MINLEN:36) and eliminating the Illumina adapters if any. Clean reads were assembled with Metaspades^65^ v3.13.1 with default parameters and k-mer iteration cycles of “21,25,31,35,41,45,51,55”. Gene annotation was performed with Prokka^66^ v1.14.5 in metagenome mode (contigs >200 bp). We assigned coding sequences to PFAMs (Pfam-A database v3.1b2) with HMMER hmmsearch v3.2.1 with the trust cut-off threshold. To determine the phylogenetic diversity of microbial communities, we used a set of 15 ribosomal proteins highly conserved across the three cellular domains, including the reduced and often fast-evolving members of the Patescibacteria and the DPANN archaea. These universal single-copy genes (USCGs; Supplementary Fig.1) were identified via their PFAM motifs (Pfam-A database v3.1b2)^67^. These were then blasted (blastp) against a collection of the 15 selected markers from all GTDB representative genomes (r214; https://gtdb.ecogenomic.org/) and assigned to the corresponding GTDB taxa when best hits had more than 35% identity over at least 70% of query lengths. To determine the relative abundance of the identified taxa, their USCGs were indexed with Bowtie2^68^ v2.3.5.1 and the clean reads from their corresponding metagenome were mapped back onto them. Mapped reads were retrieved with Samtools^69^ v1.9 and, for each gene, the Reads Per Kilobase (of gene sequence) per Million of mapped metagenomic reads (RPKMs) were calculated using *ad hoc* Perl scripts, allowing normalization for gene length and sample sequencing depth. We then averaged the results from the 15 USCGs. Plots were generated with ad hoc R scripts using the ggplot2 package^70^ at the phyla level (Fig.1) and the genus level for haloarchaea and Nanohaloarchaea (Extended Data Fig.1).

### Diversity indexes

To calculate diversity indexes, we used operational taxonomic units (OTUs) defined by clusters of USCG genes at 99% identity. Clusters were generated by cd-hit^71^ (parameters -c 0.99, -n 5, -d 0, -M 0) and assigned to species-level taxa (GTDB r214) as described above. Species abundance was calculated by the sum of RPKMs of clustered genes for each OTU after averaging the RPKMs for the different USCGs assigned to the same species. These values served to build an abundance matrix and calculate several diversity indices (alpha diversity, Shannon entropy, Pielou’s evenness and Simpson’s dominance) using an *ad hoc* R script using the Vegan package^72^. These were visualized as lollipop plots using the ggplot2 R package. In addition to diversity indexes for the Danakil metagenomes, we included data from metagenomes of freshwater, Mediterranean plankton and solar saltern ponds previously generated and treated in the same way^35^ for comparison.

### Metagenome-assembled genomes (MAGs)

To generate MAGs, we co-assembled reads from, respectively, Ass and 9Ass (co-assembly DAL-Ass), and WCL2 and WCL3 (co-assembly DAL-WCL) with Metaspades using the same parameters as above (statistics shown in Supplementary Table 1). These co-assemblies were used for binning via the anvi’o pipeline^73^ v5, using Concoct^74^ v1.1.0 and Metabat^75^ v2.15 as binning software and the DASTool^76^ v1.1.2 to merge and dereplicate bins. The resulting bins were manually refined using the anvi-interactive graphical user interface to obtain high quality MAGs. In the case of the single 9Gt metagenome assembly, MAGs were binned using Concoct using default parameters except for contig splitting (10,000 nt fragments). Low-yield metagenomes from La Grotte were mapped onto the 9Gt Concoct bins using anvi’o to enable the use of the anvi-interactive graphical user interface for manual bin refinement. Final MAG identifiers (IDs) consist of the prefix DAL-(Dallol region), the name of the metagenome where they binned from (DAL-9Gt, DAL-Ass, DAL-WCL), followed by the original bin number and a label indicating completion and redundancy as determined by anvi’o (e.g. 90C3R indicates 90% completion and 3% redundancy). Completion and redundancy values were independently inferred using CheckM^77^ v1.1.1 and CheckM2 v1.0.1. We assembled a total of 483 MAGs, from which 155 had good quality (>40% completion and <5% redundancy; Supplementary Table 3). MAGs were annotated with prokka^66^ specifying the domain of each respective MAG (Archaea or Bacteria). Coverage was only calculated for the set of good-quality MAGs, and it was determined with CoverM^78^ v0.6.1 by mapping the corresponding metagenomic reads to each of the MAG contigs. Different MAG statistics are shown in Supplementary Table 3.

### Phylogenetic classification of MAGs

All assembled MAGs were initially classified using the GTDB-tk pipeline^37^ v2.3.0 according to the GTDB r214 taxonomy. Average nucleotide identity (ANI) comparisons among groups of MAGs were conducted using the ANI matrix calculator^79^. High-quality MAGs were more robustly classified according to their phylogenetic position in phylogenomic analysis; their statistics are shown in Supplementary Table 3. The phylogenomic analyses for archaea were carried out using a subset of 127 proteins from a previously identified set of core archaeal markers allowing to robustly infer the tree of archaea^3,80^. We added our Danakil MAGs to an initial sample of 178 proteomes spanning all major archaeal super-groups^3^ and additional GTDB representative genomes for the Halobacteriota, Aenigmatarchaeota, and Nanohaloarchaeota. We gathered orthologs of the 127 markers from the Danakil MAG predicted proteomes by using sequences from previous alignments^3^ as queries for BLASTp. For each marker, the best BLAST hit from each proteome was added to the dataset. Each dataset was aligned using mafft-linsi ^81^ and ambiguously aligned positions were trimmed using TrimAl^82^ (‘automatic1’ mode). All trimmed alignments were concatenated into a supermatrix (533 taxa, 32,985 amino acid positions). A preliminary phylogeny was inferred using FastTree^83^ v2 (-lg -gamma model of evolution) that served as basis to select two subsets of taxa focusing on the DPANN and the Halobacteriota (188 and 260 taxa, respectively). Those subsets were used for phylogenetic reconstruction with IQ-TREE v2^84^ under the LG + C60 + G model; 1,000 ultrafast bootstrap replicates were used to assess branch statistical support. Independent phylogenomic trees were carried out for the different bacterial clades detected (T1Sed10-126, Cyanobacteria, Myxococcota, Rhodothermia, including Salinibacteraceae, Patescibacteria, and Bipolaricaulota). Representative reference genomes were selected from GTDB for each group, and the 120 gene markers used by the GTDB-tk classifier retrieved. The corresponding proteins were aligned with mafft^85^ v7.453, and ambiguously aligned positions trimmed with trimAl^82^ v1.4.rev22 prior to their concatenation with an *ad hoc* Perl script. ML phylogenetic trees were reconstructed with IQ-tree^86^ v1.6.11 with the following parameters: -bb 1000 -m LG+C60+G+F. Full trees for Fig.4 are provided as Supplementary Figs.4-6; other bacterial trees are shown in Supplementary Figs.7-11.

### Isoelectric point, amino acid biases and statistical analyses

To calculate the pI of each protein from metagenomes and MAGs, we used the EMBOSS’ iep software^87^. The output was processed with an *ad hoc* Perl script, and density plots were visualized using an *ad hoc* R script with the ggplot2 package. Median pI values for each proteome were calculated using a Perl script. Amino acid ratios (R/K and D+E/I+K) were calculated for each inferred proteome with an *ad hoc* Perl script (AAfreq.pl). Metagenome R and K frequency bar-plots were built with an *ad hoc* R script using the ggplot2 package. pI and amino acid frequencies and ratios are shown in Supplementary Table 2. Principal Component Analyses (PCA) including those values as well as some physicochemical parameters (Supplementary Table 2) were carried out with the ggbiplot and corrplot packages in R^88^.

### Metabolic inference

The general metabolic potential of MAGs with more than 50% completion was automatically evaluated using the Metabolic-G software^89^ v4.0. The resulting tables 2 (Function Hit) and 3 (KEGG Module Hit) were used for visualization via an *ad hoc* R script. Function Hit was visualized as a heatmap with the presence/absence of each function using the gplots package. For the KEGG module Hit, pathway completeness was calculated based on the number of steps needed to achieve the module function. A gradient of 0-100% completion of the module was visualized via heatmap with the gplots R package. The haloarchaeal and nanohaloarchaeal MAG heatmaps were ordered according to guiding phylogenetic trees using the phylogram R package. To look for diagnostic genes, we used the collection of HMMs from KOfams developed by KEGG^90^ to scan the whole metagenomic space with hmmsearch (e-value <1e-^11^). To look for more specific functions, we predicted and annotated genes following adapted pipelines^91^; marker genes for reduction and oxidation of both organic and inorganic compounds were searched manually. We annotated the genes against the UniProtKB-SwissProt (05.2022) using Diamond^92^ blastp v2 (parameters: -k1 --evalue 1e-10 --query-cover 50 --id 40 --sensitive) and the Pfam database (release 35.0) using HMMsearch v3.3.2 (parameters: cut_ga). Carbohydrate-active enzymes (CAZymes) were annotated using the specialized database CAZyDB (release 09242021). The number of genes was normalized based on genome estimated number, calculated by average count of USCG and sequence depth following MicrobeCensus^93^ v1.1.1. The metabolic potential of the most complete MAGs was used to illustrate the metabolism of given clades (Fig.5; Supplementary Table 4).

## Data availability

Metagenomes are available in GenBank with the following accession numbers under the Bioproject PRJNA541281: SAMN37693137 (DAL-Ass), SAMN37693138 (DAL-9Ass), SAMN37693139 (DAL-9Gt), SAMN37693140 (DAL-WCL2) and SAMN37693141 (DAL-WCL3).

## Code availability

Custom code for these analyses (*perl* and R scripts) is available at Gitlab https://gitlab.com/DeemTeam/dal-metagenomes.

## Acknowledgements

We thank F. Brenckman and the Iris Foundation for supporting our initial field trip in 2016 and the Mamont Foundation, for field trip support in 2019. We thank J. Belilla, J.M. López-García, A.I. López-Archilla, K. Benzerara, L. Jardillier, O. Grunewald, L Cantamessa and the Afar authorities for their assistance during field trips and P. Deschamps for help with software installation. This work was supported by the Moore-Simons Project on the Origin of the Eukaryotic Cell (P.L.-G., https://doi.org/10.37807/GBMF9739), the Iris Foundation (https://en.fondationiris.org/) and the European Research Council Advanced Grant Plast-Evol (D.M., No. 787904).

## Author contributions

P.L.-G. and D.M. organized the field trips, collected and conditioned samples, designed the research and obtained funding to conduct it. A.G.-P. carried out the bioinformatic, phylogenetic and statistical analyses from raw metagenome data. B.D. and A.G.-P. investigated genes involved in metabolism. B.B. and L. Eme carried out the final phylogenomic analysis for Halobacteria and Nanohaloarchaeota. P.L.-G. conceptualized and supervised the research, and wrote the manuscript. All authors read and commented on the manuscript.

---

**Extended Data Fig. 1.**
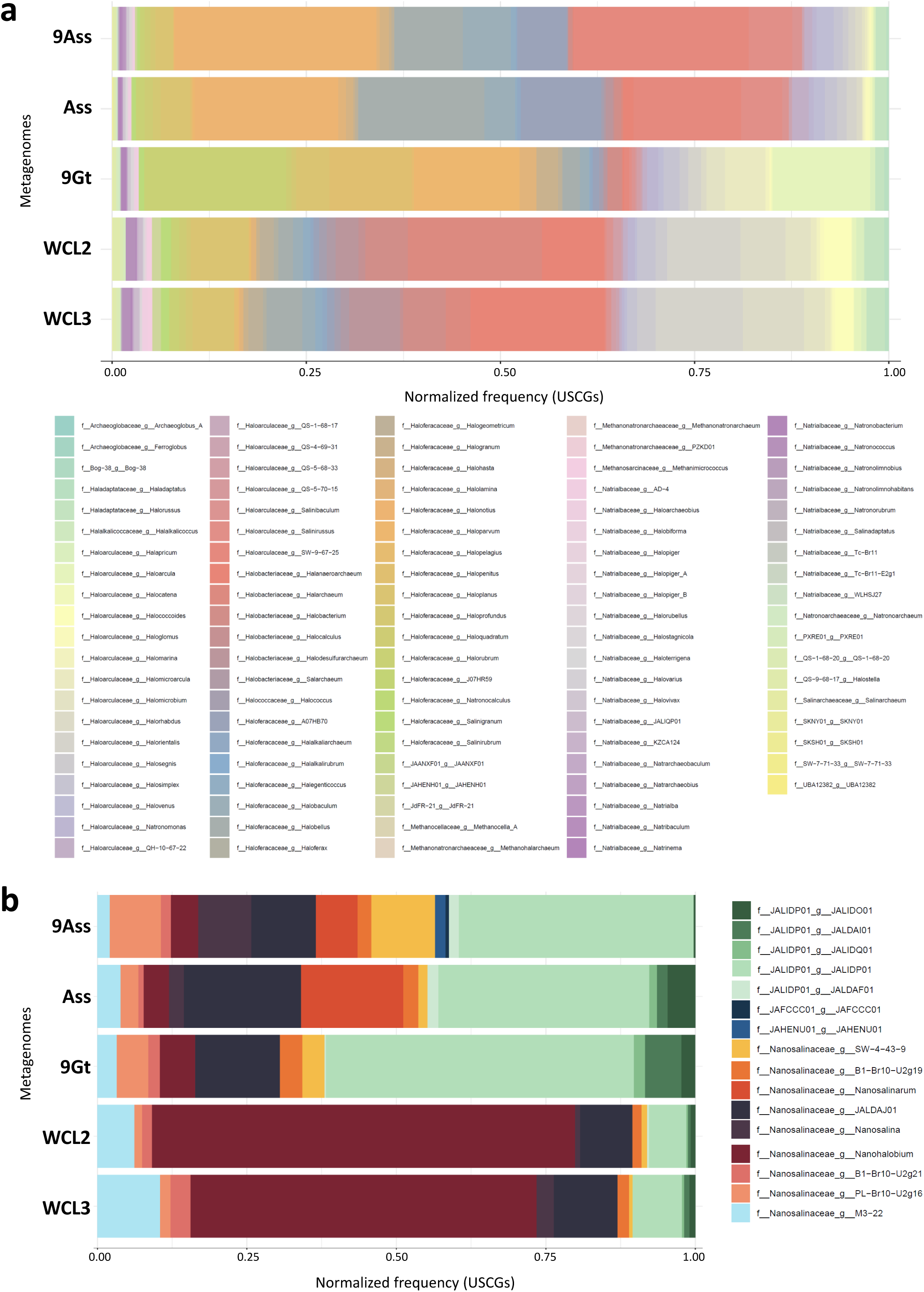
Genus-level composition of dominant archaeal phyla in polyextreme ecosystems in the north Danakil Depression. **a**, Composition of members ascribing to the Halobacteriota. **b**, Composition of members ascribing to the Nanohaloarchaeota. Classification according to GTDB r214.

**Extended Data Fig. 2.**
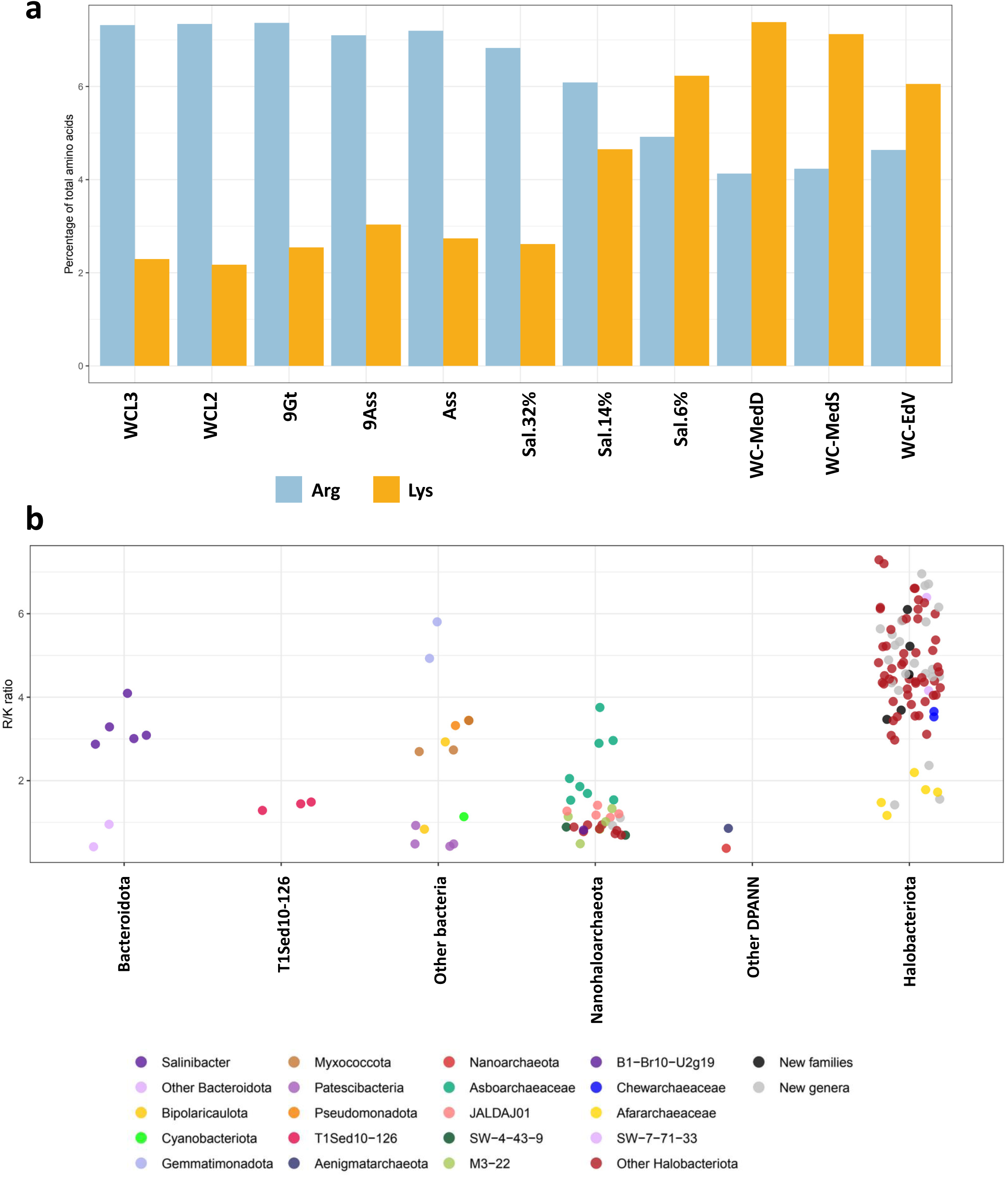
Arginine preference over lysine in north Danakil extreme halophilic communities. **a**, Percentage of arginine and lysine in proteomes inferred from Danakil polyextreme ecosystems in comparison with representative ecosystems along a salinity gradient. **b**, Jitter plot showing Arg/Lys (R/K) ratio inferred from MAGs of Danakil chaotropic brines classified by major taxa.

**Extended Data Fig. 3.**
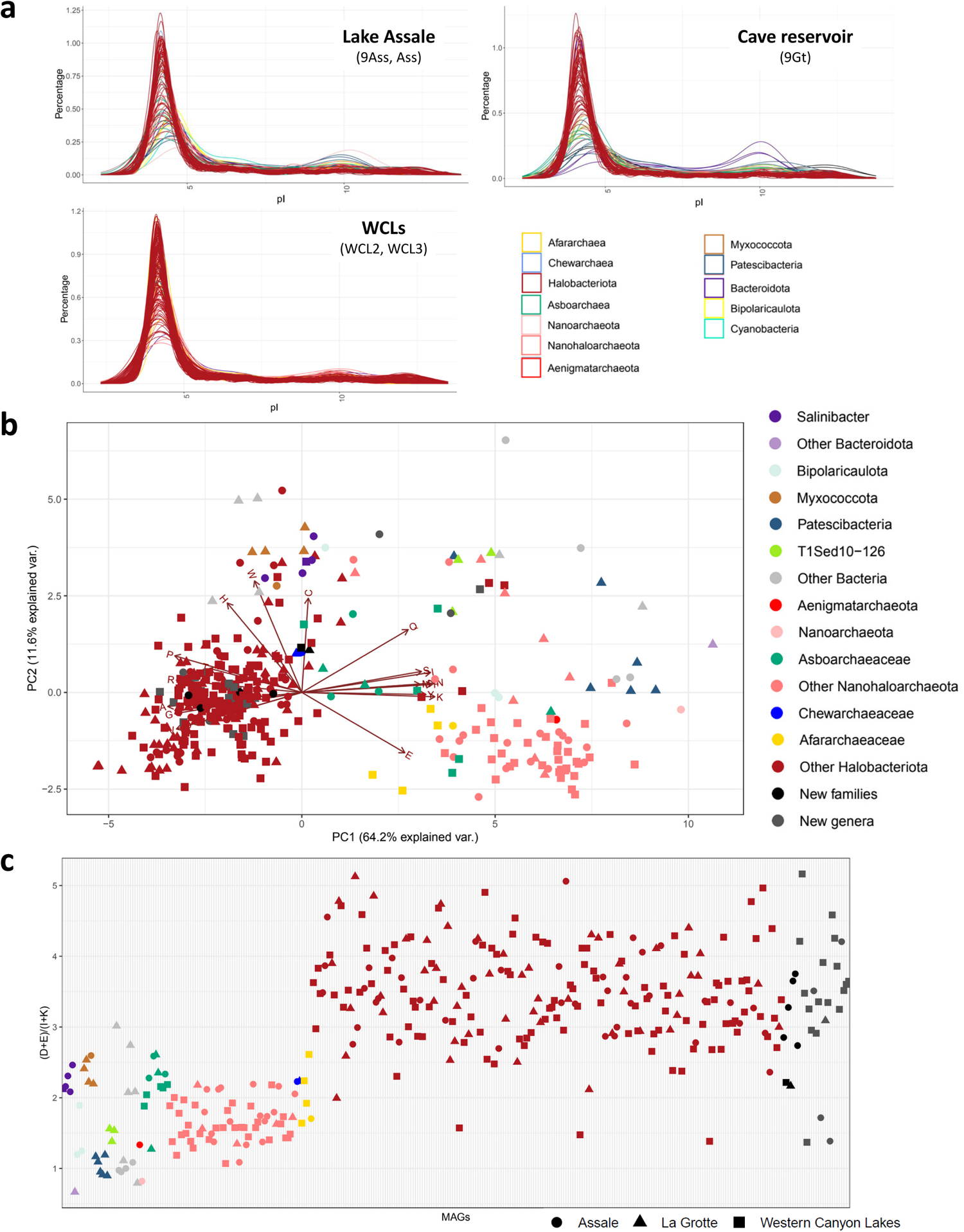
Isoelectric point (pI) and amino acid biases in MAGs assembled from polyextreme brines in the north Danakil Depression. **a**, Distribution of pI values in individual MAGs assembled from Lake Assale (Ass, 9Ass), cave reservoir La Grotte (9Gt) and Western Canyon Lakes (WCLs) metagenomes. **b**, PCA of amino acid composition and individual MAGs retrieved from polyextreme north Danakil ecosystems. **c**, Asp-Glu/Ile-Lys (DE/IK) ratio in the assembled MAGs. MAGs are colored according to their taxonomic affiliation. Symbol shapes denote the type of hypersaline ecosystem the MAGs were assembled from.

**Extended Data Fig. 4.**
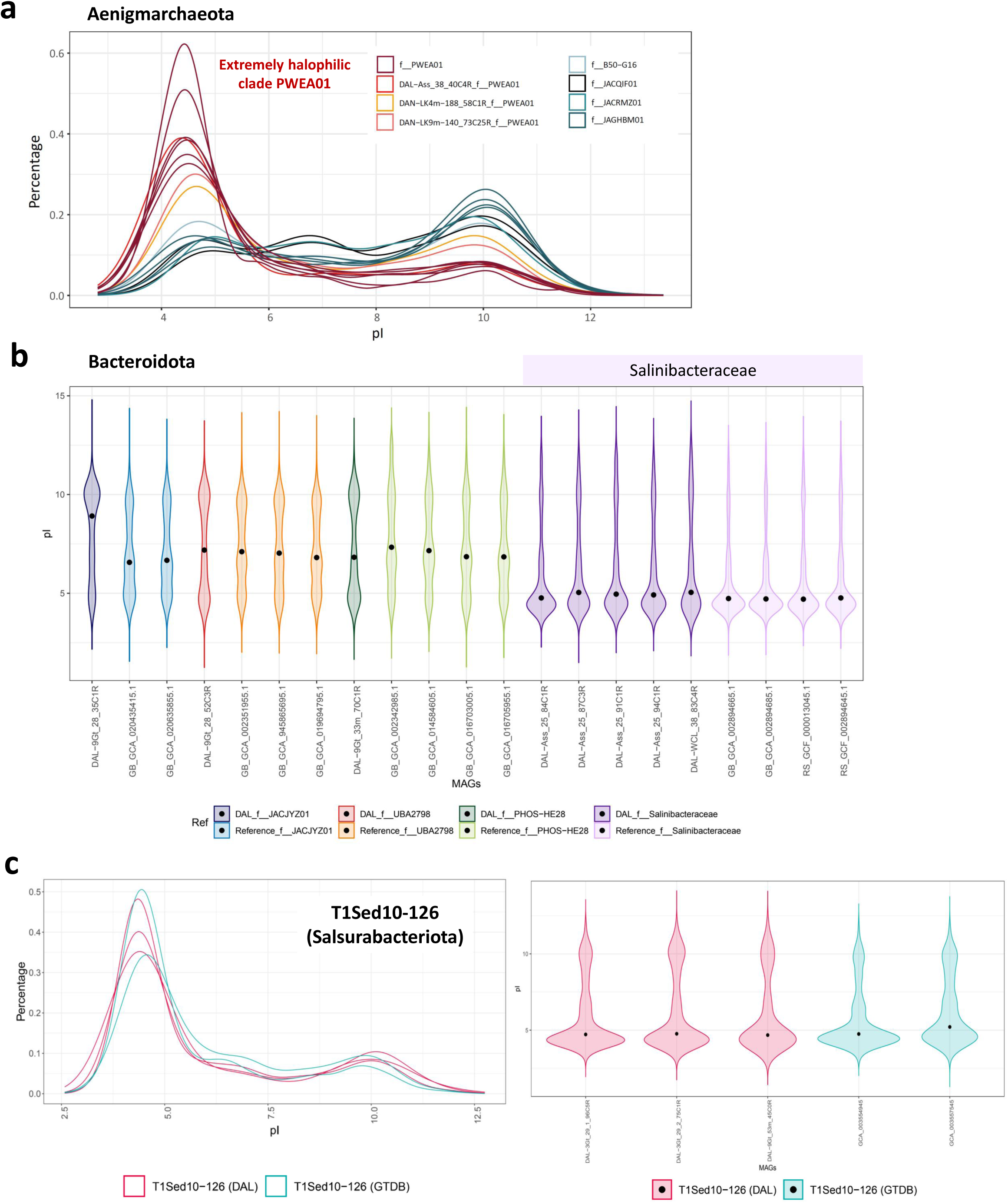
Isolelectric point of proteomes inferred for archaeal and bacterial lineages present in north Danakil hypersaline systems independently adapted to hypersaline conditions. **a**, Distribution of pI values in individual MAGs belonging to the Aenigmarchaeota order PWEA01, including the Dallol area MAGs (reddish color), and its phylogenetic relatives (blue colors) as reference. **b**, Violin plots showing pI values for Danakil MAGs belonging to the Bacteroidota, including references for comparison. Note the low pI values characteristic of the extremely halophilic Salinibacteraceae. **c**, Distribution of pI values shown by curves (left) and violin plot (right) of MAGs ascribing to the phylum T1Sed10-126 (*Candidatus* Salsurabacteriota), including the three genomes assembled from the Dallol area ecosystems and the two existing reference genomes from GTDB.

**Extended Data Fig. 5.**
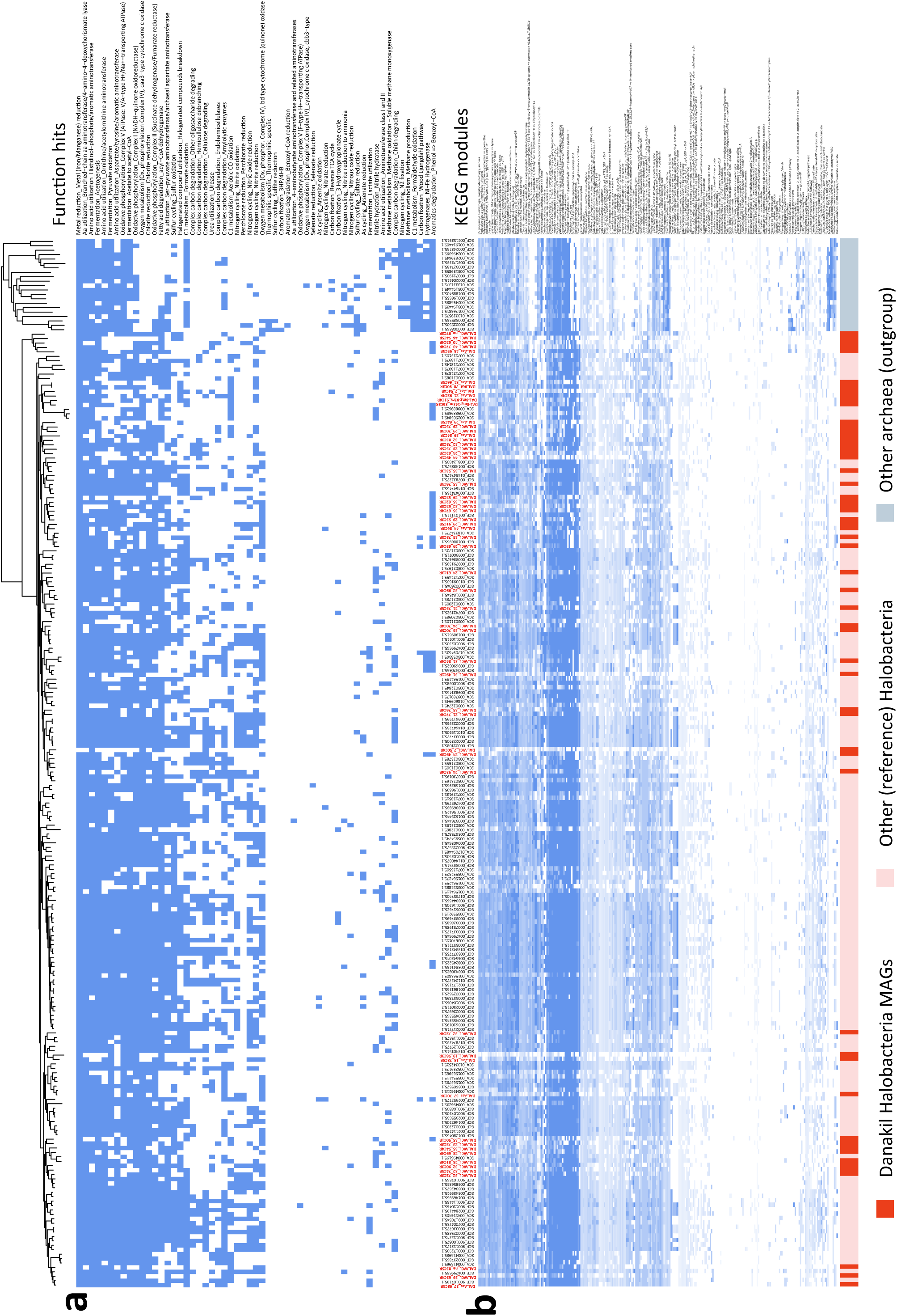
Heat maps showing major metabolic potential functions and KEGG pathways for individual Halobacteria MAGs assembled from north Danakil hypersaline ecosystems. **a**, Presence (blue)-absence (white) of metabolic functions as inferred by Metabolic G in Danakil MAGs and representative GTDB genomes of Halobacteria. **b**, Modules of KEGG pathways identified in Halobacteria MAGs and GTDB reference genomes (outgroup). The blue color intensity indicates the completeness of the respective pathway (intense, complete). The names of our MAGs are highlighted in red. Aa, amino acid.

**Extended Data Fig. 6.**
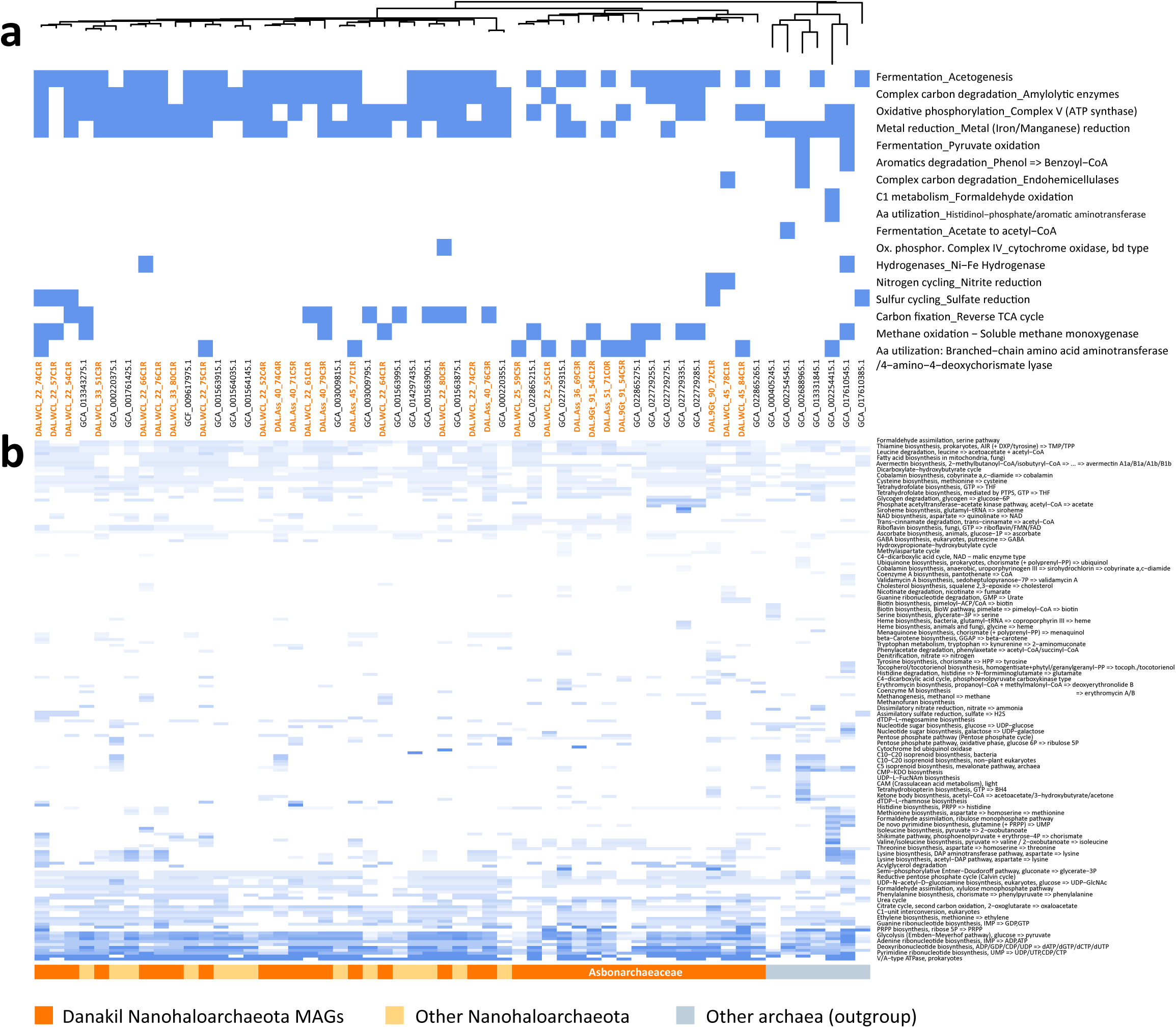
Heat maps showing major metabolic potential functions and KEGG pathways for individual Nanohalobacteriota MAGs assembled from north Danakil hypersaline ecosystems. **a,** Presence (blue)-absence (white) of metabolic functions as inferred by Metabolic G in Danakil MAGs and representative GTDB genomes of Nanohaloarchaeota. **b**, Modules of KEGG pathways identified in Nanohaloarchaeota MAGs and GTDB reference genomes (outgroup). The blue color intensity indicates the completeness of the pathway. The names of our MAGs are highlighted in orange. Aa, amino acid; ox. phosphor.

**Extended Data Fig. 7.**
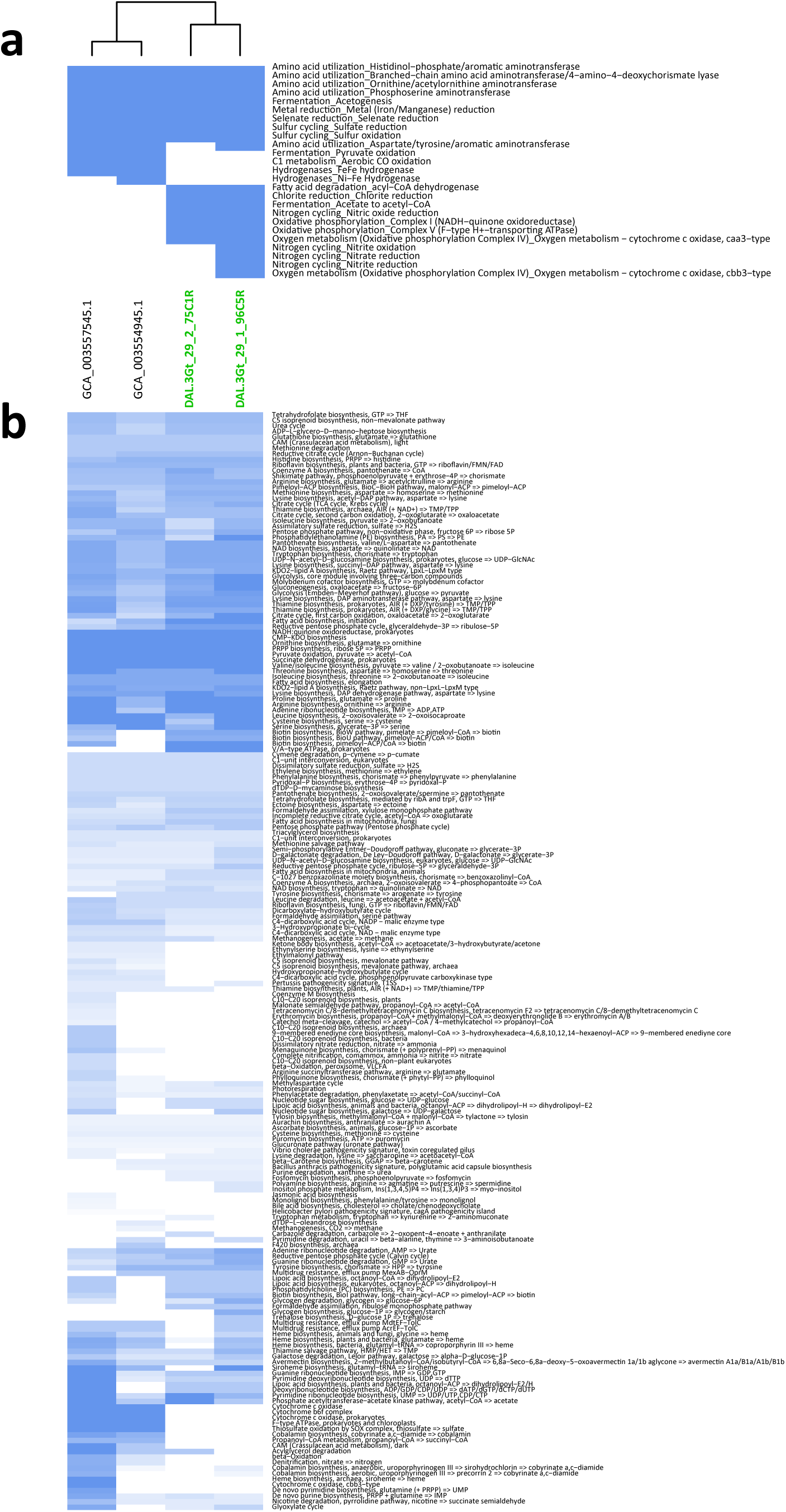
Heat maps showing major metabolic potential functions and KEGG pathways for MAGs of the halophilic candidate bacterial phylum Salsurabacteriota (T1Sed10-126). **a,** Presence-absence of metabolic functions as inferred by Metabolic G in the two most complete Danakil MAGs and known GTDB genomes. **b**, Modules of KEGG pathways identified in Salsurabacteriota. The names of our MAGs are highlighted in green.

**Extended Data Fig. 8.**
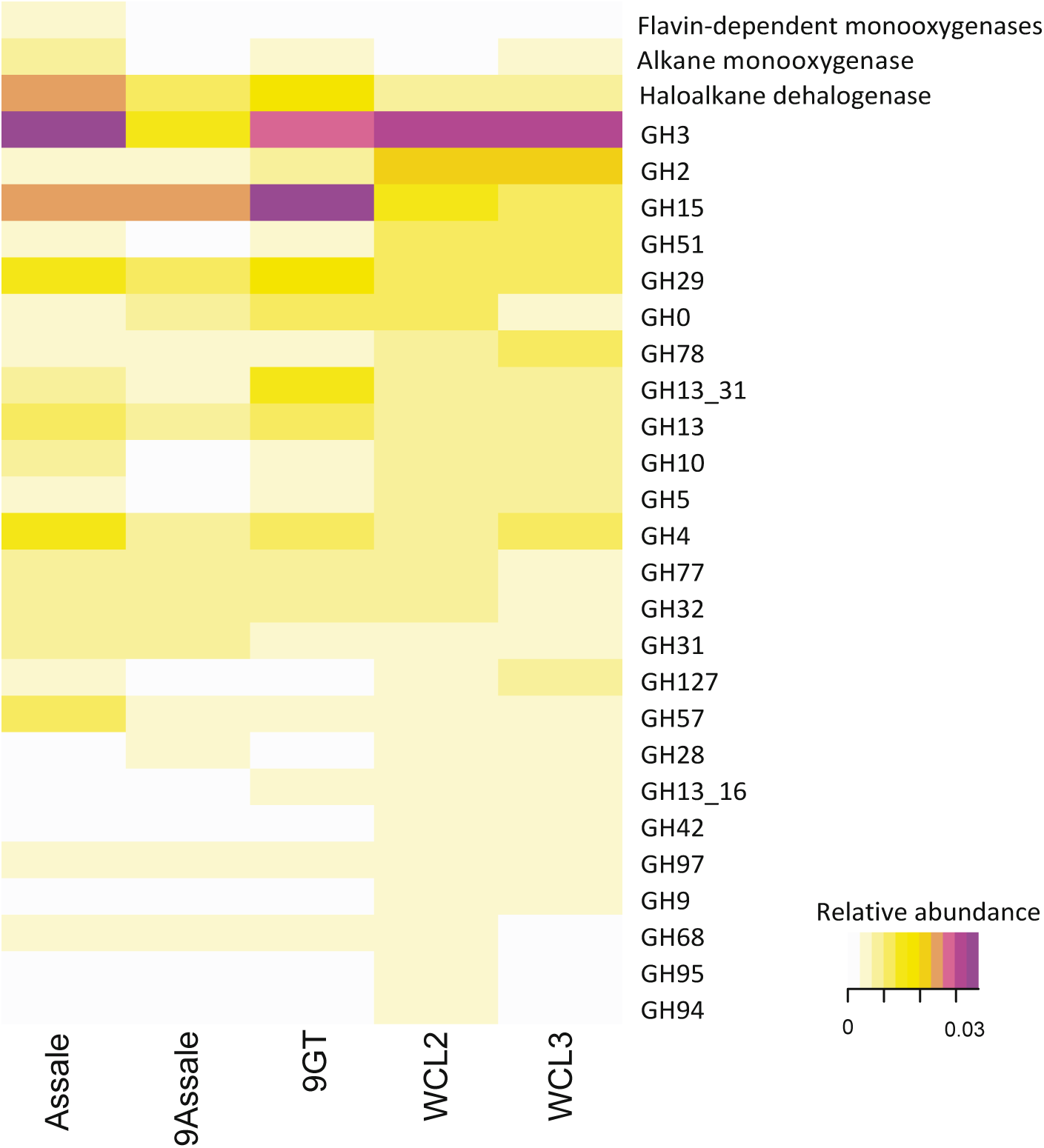
Heat map showing the relative abundance of hydrocarbon degradation genes in metagenomes from north Danakil hypersaline ecosystems.

