## Supplementary Figures 1-11 for "Extremely acidic proteomes and metabolic flexibility in bacteria and highly diversified archaea thriving in geothermal chaotropic brines"

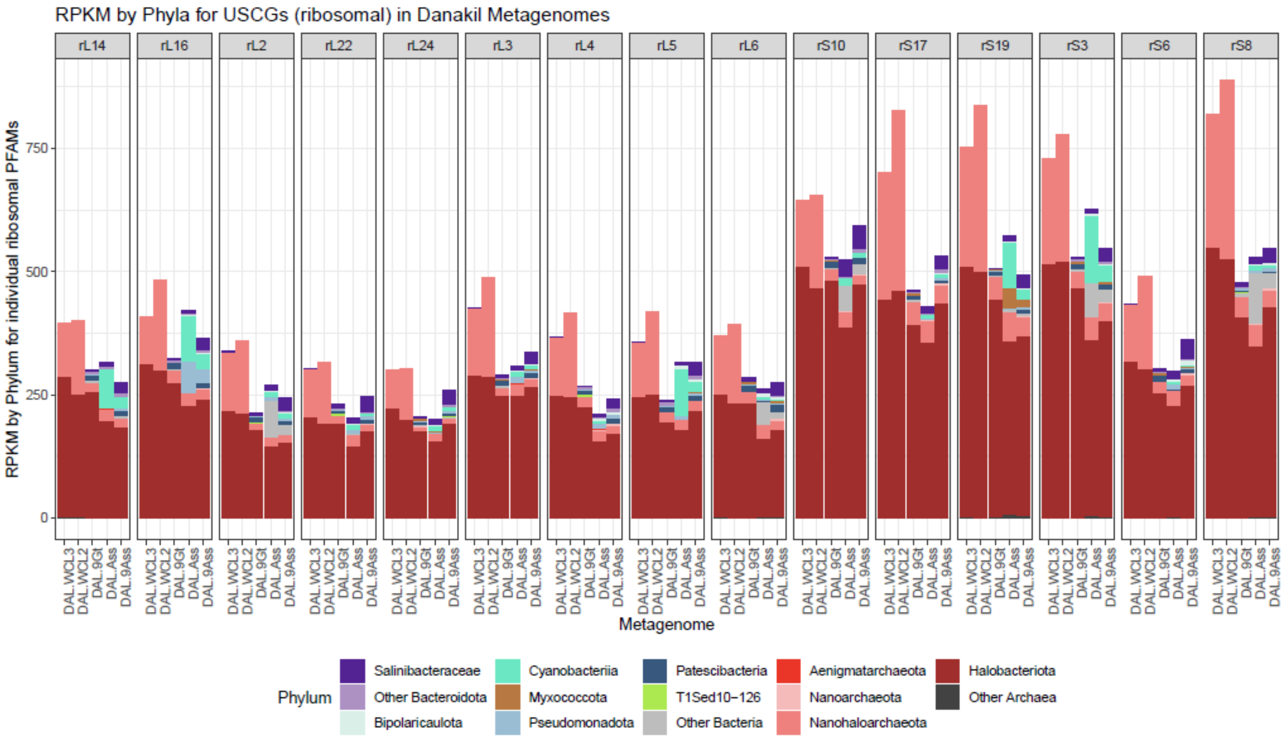


**Supplementary Fig. 1. Normalized abundance of selected ribosomal proteins used as universal single-copy genes assigned to microbial taxa.**

a

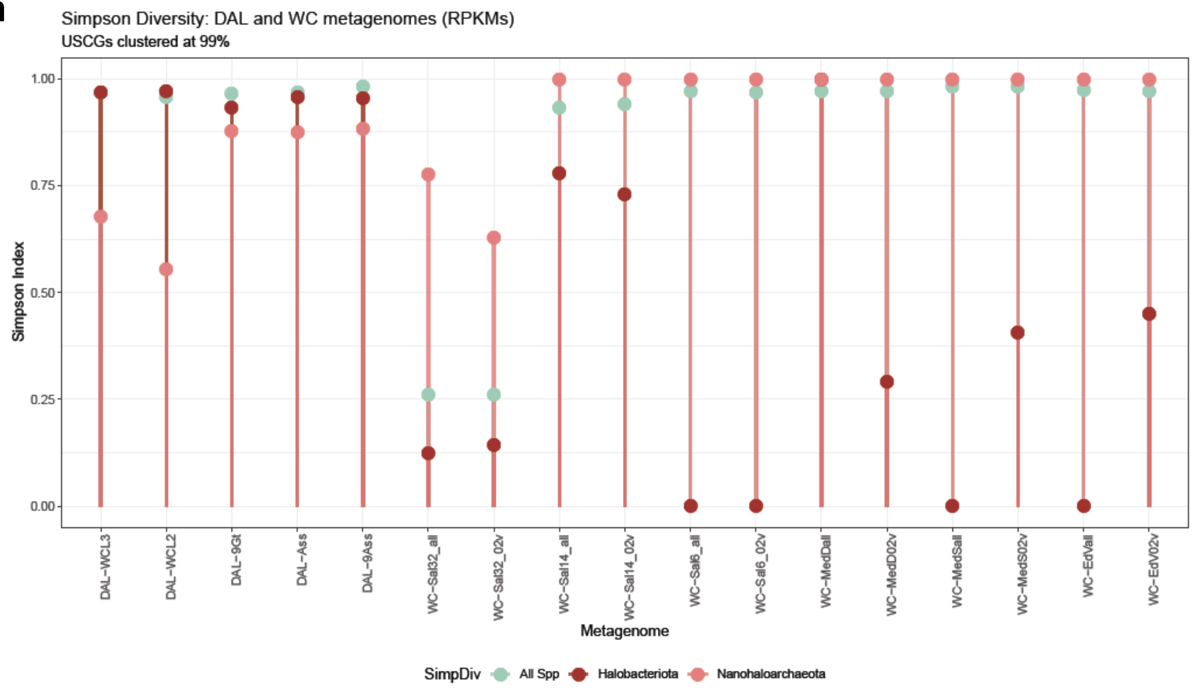


b

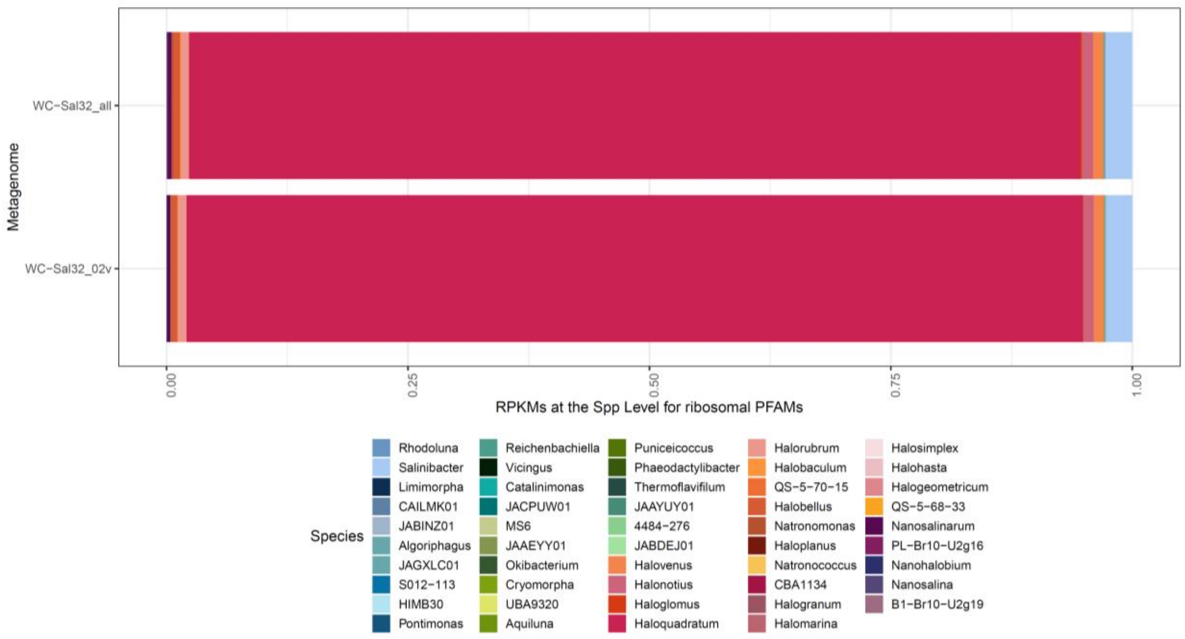


**Supplementary Fig. 2. Simpson diversity index calculated for all OTUs as well as Halobacteria and Nanohaloarchaeota (a) and species-level community composition inferred from replicate metagenomes of solar saltern 32% salinity ponds (b).**

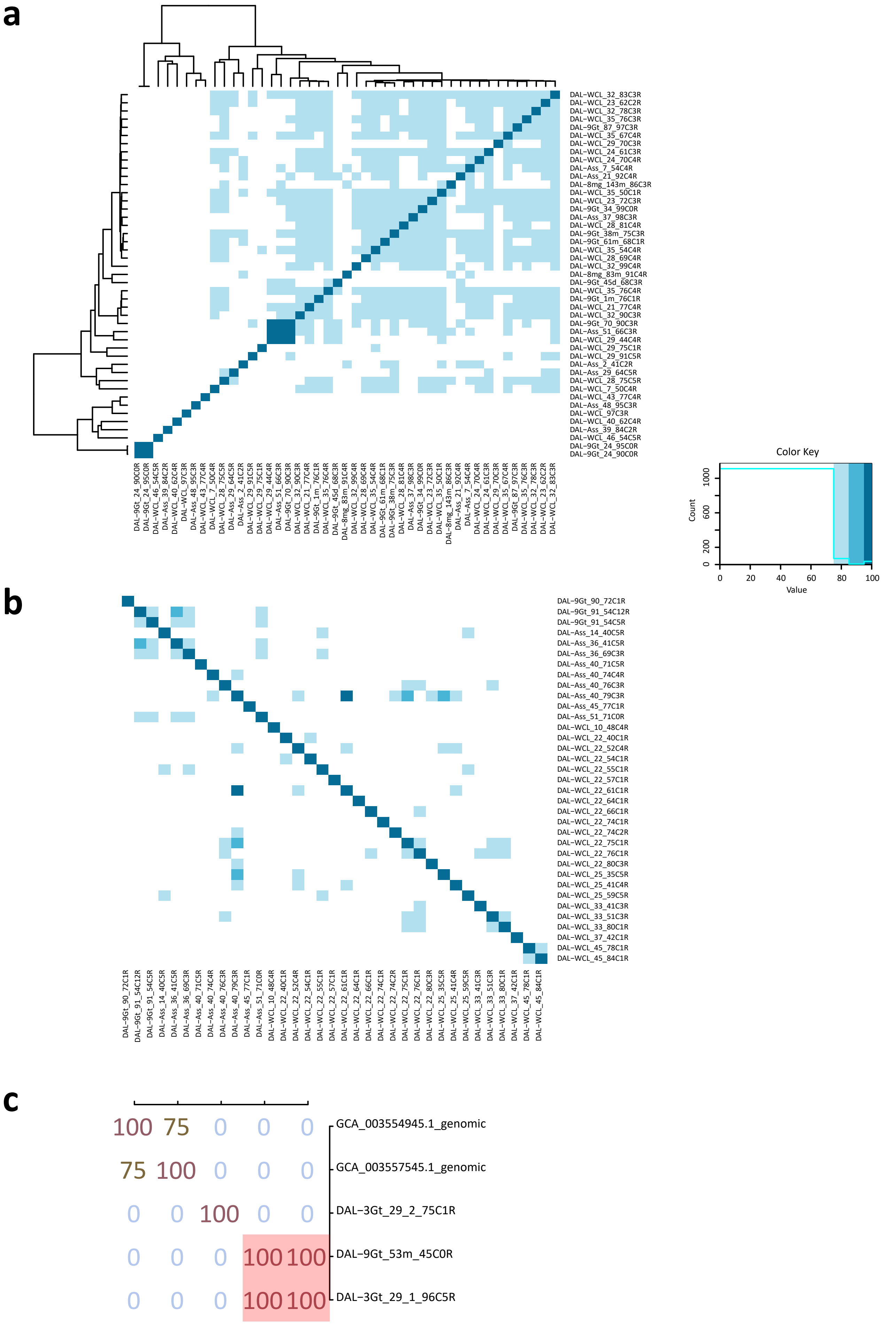


**Supplementary Fig. 3. Average nucleotide identity values among MAGs affiliating to different lineages. a,** Halobacteria (Halobacteriota). **b**, Nanohaloarchaeota. **c**, Salsurabacteriota (p_T1Sed10-126).

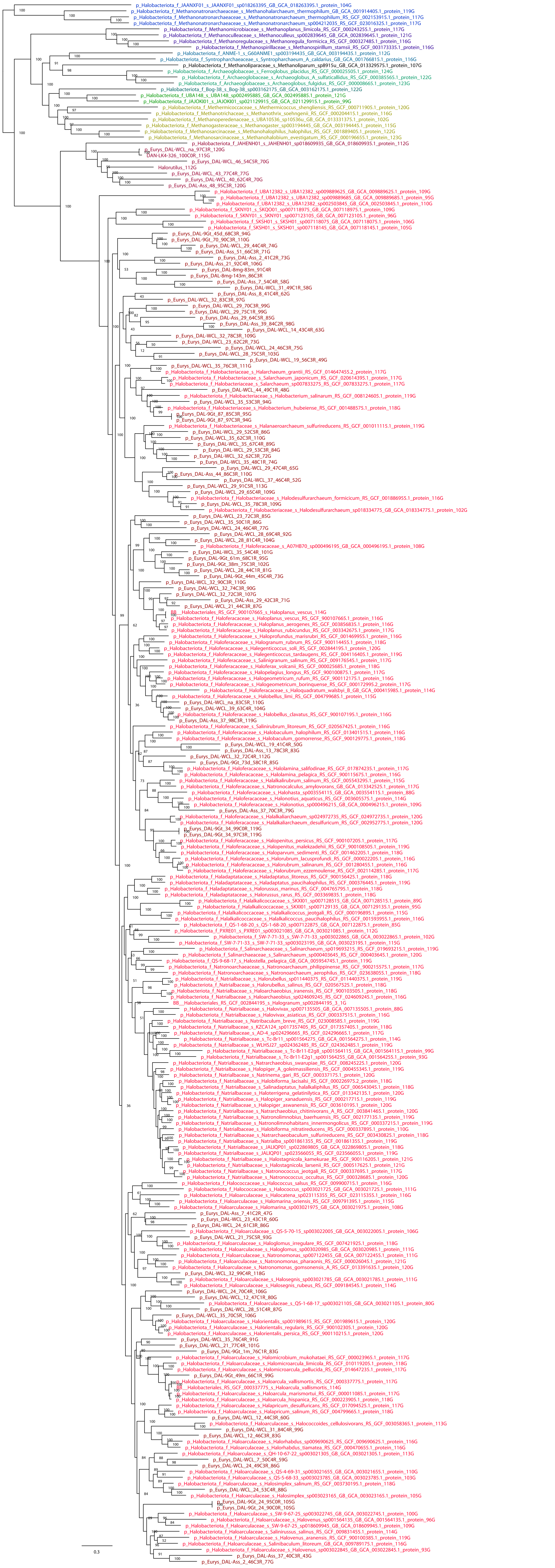

**Supplementary Fig. 4. Detailed ML phylogenomic tree of Halobacteria**

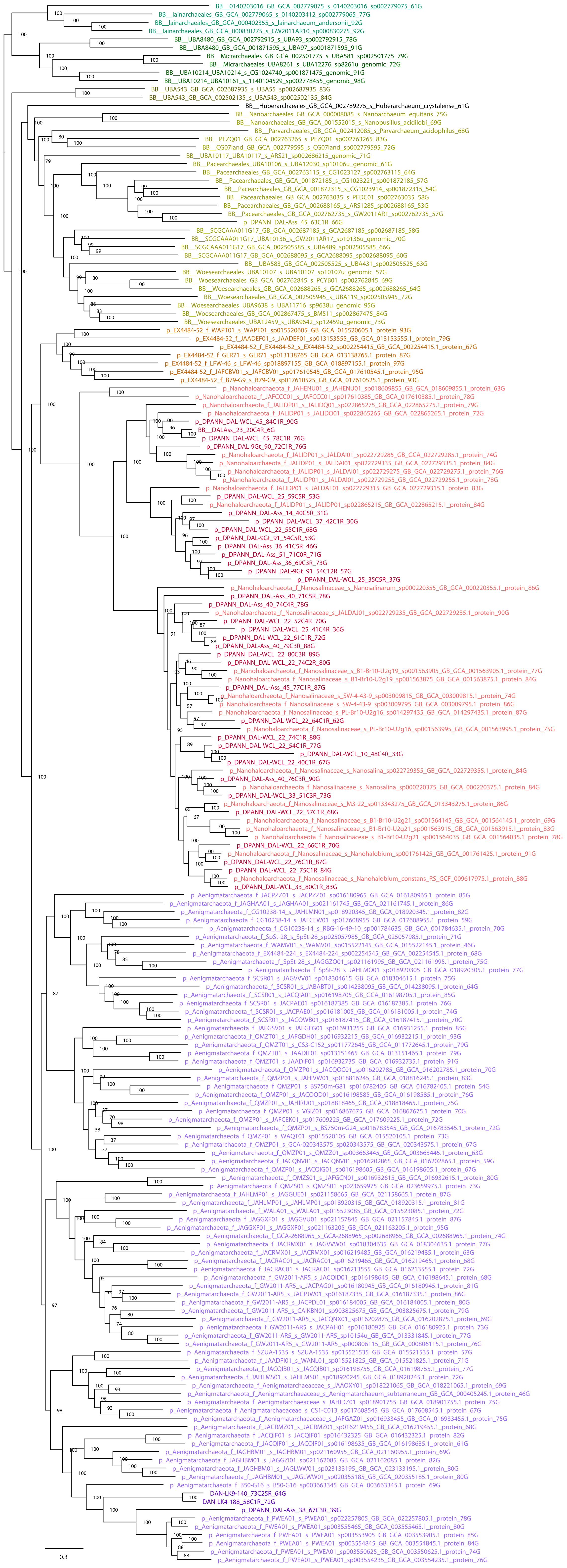


**Supplementary Fig. 5. Detailed ML phylogenomic tree of Nanohaloarchaeota**

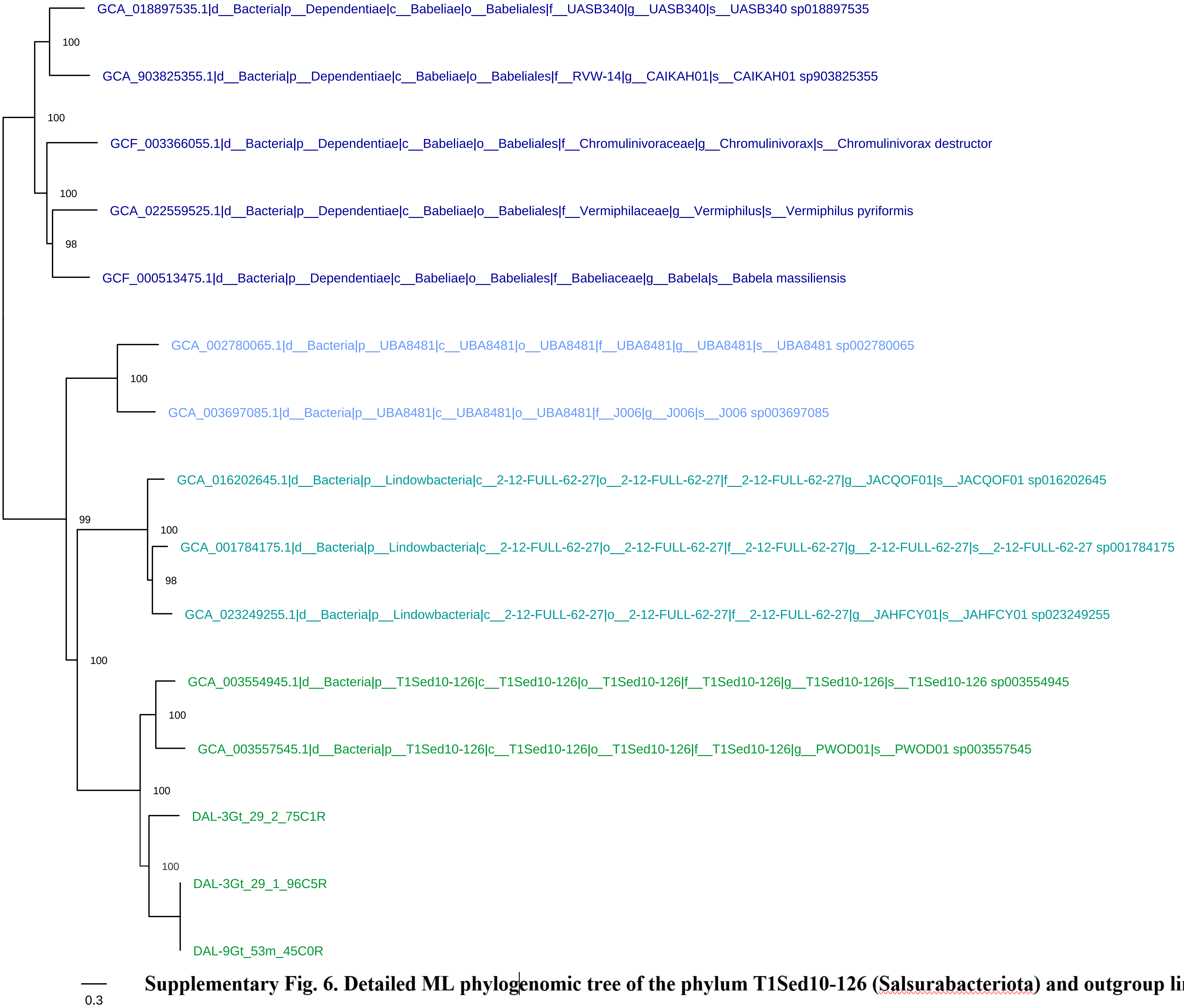


**Supplementary Fig. 6. Detailed ML phylogenomic tree of the phylum T1Sed10-126 (Salsurabacteriota) and outgroup lineages.**

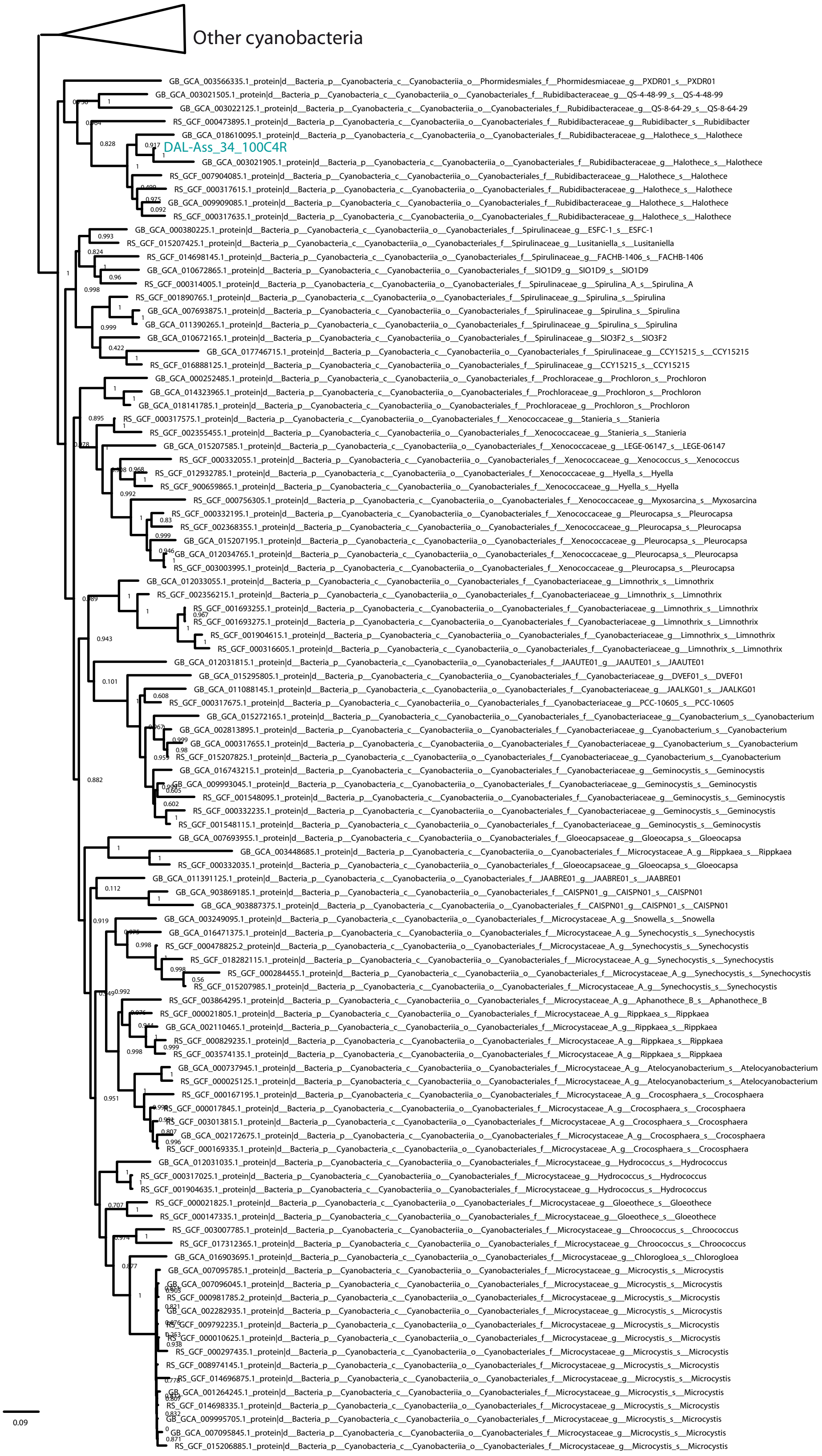


Supplementary Fig.7. Phylogenetic tree of Cyanobacteria

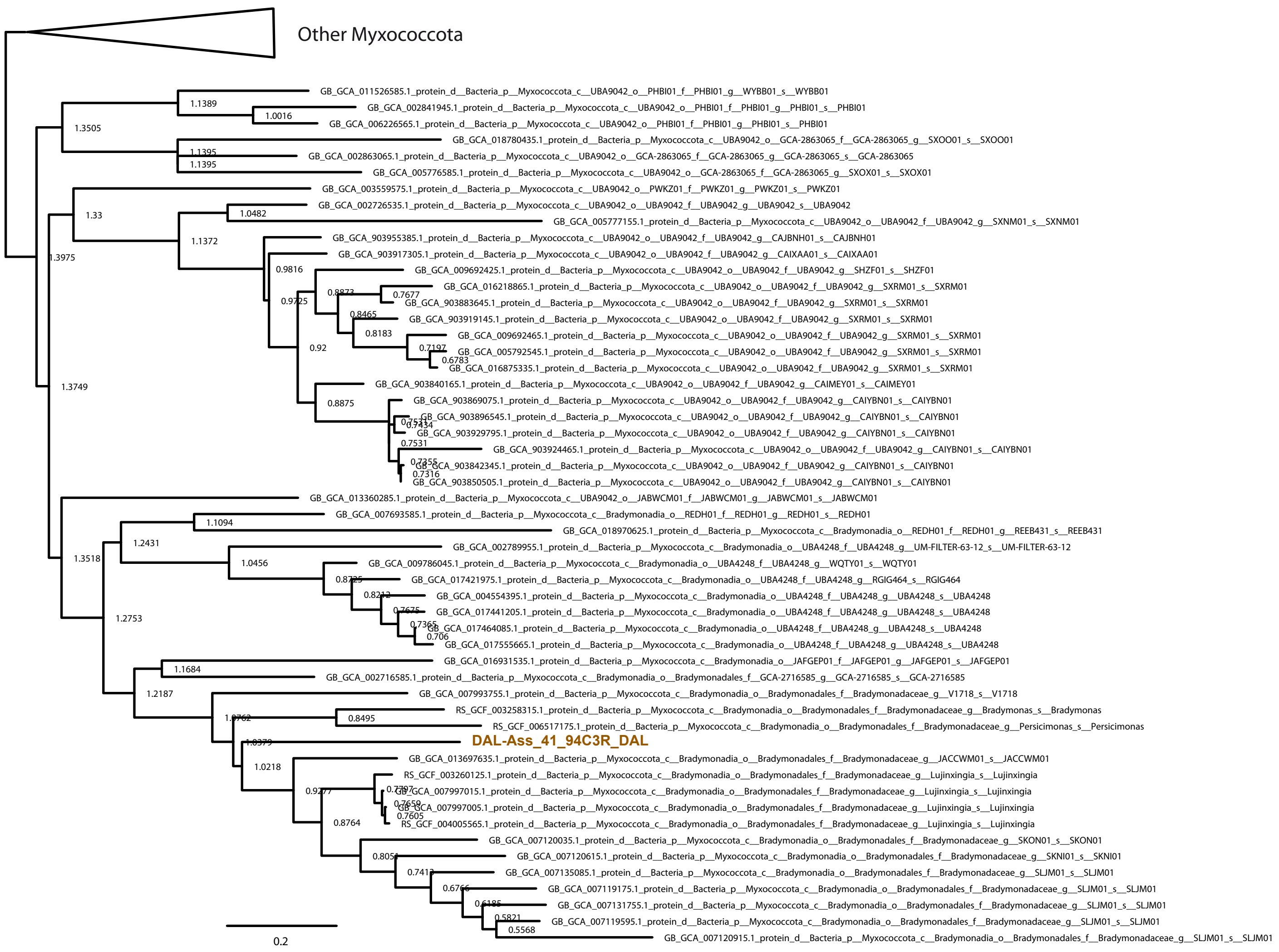


Supplementary Fig.8. Phylogenetic tree of the Danakil Myxococcota

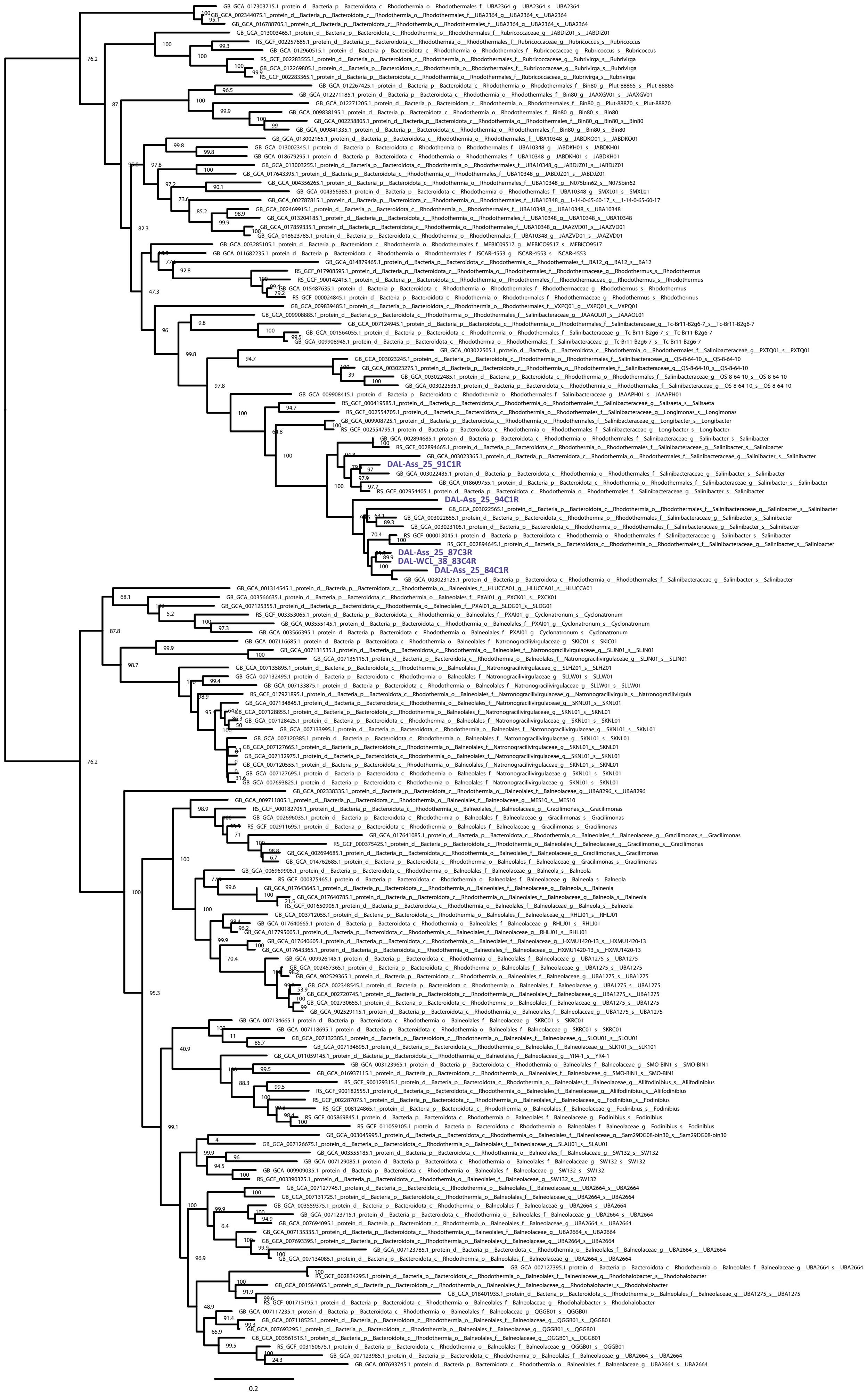


Supplementary Fig.9. Phylogenetic tree of the Salinibacteraceae (Rhodotermia)

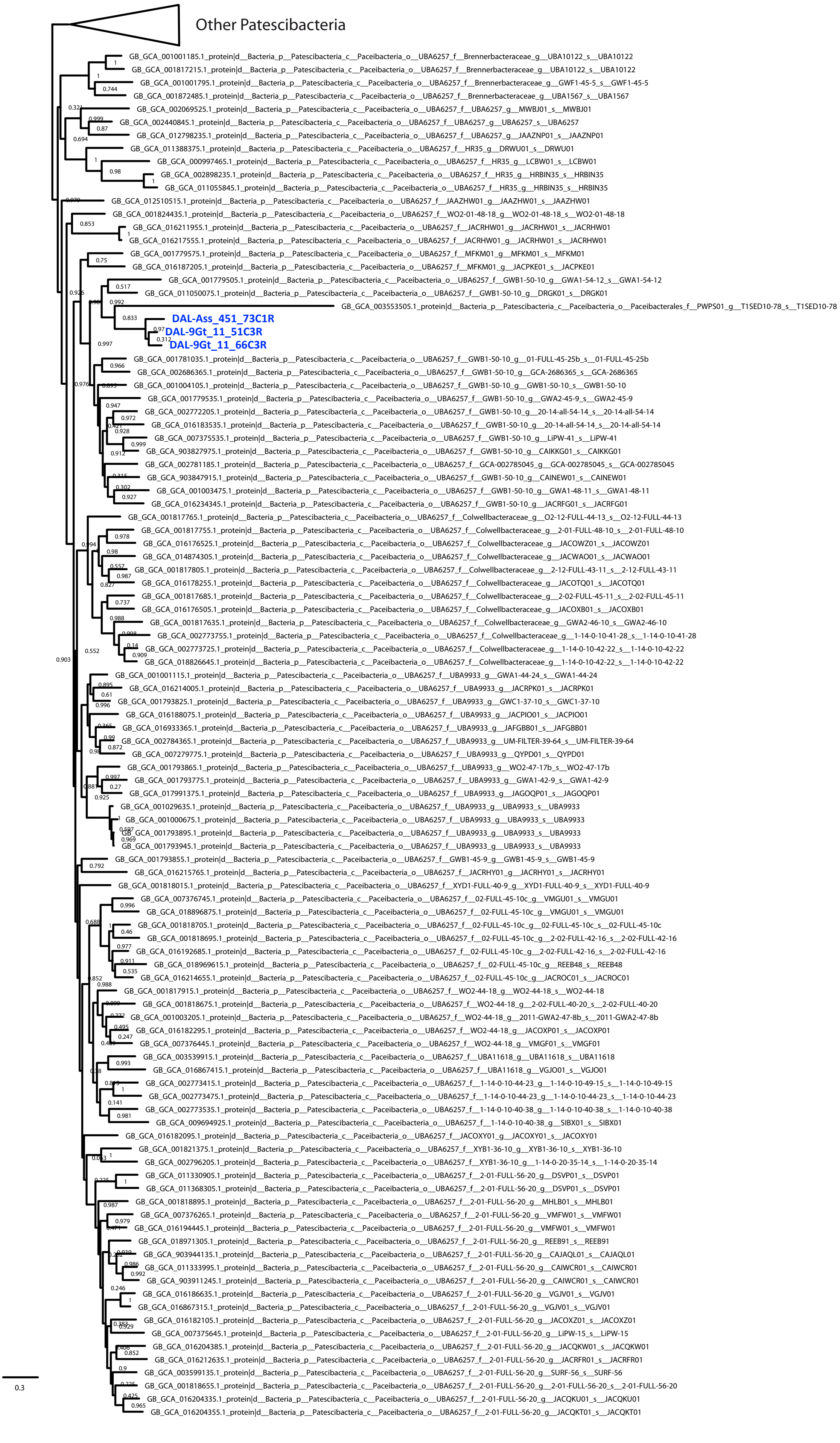


Supplementary Fig.10. Phylogenetic tree of the Patescibacteria MAGs

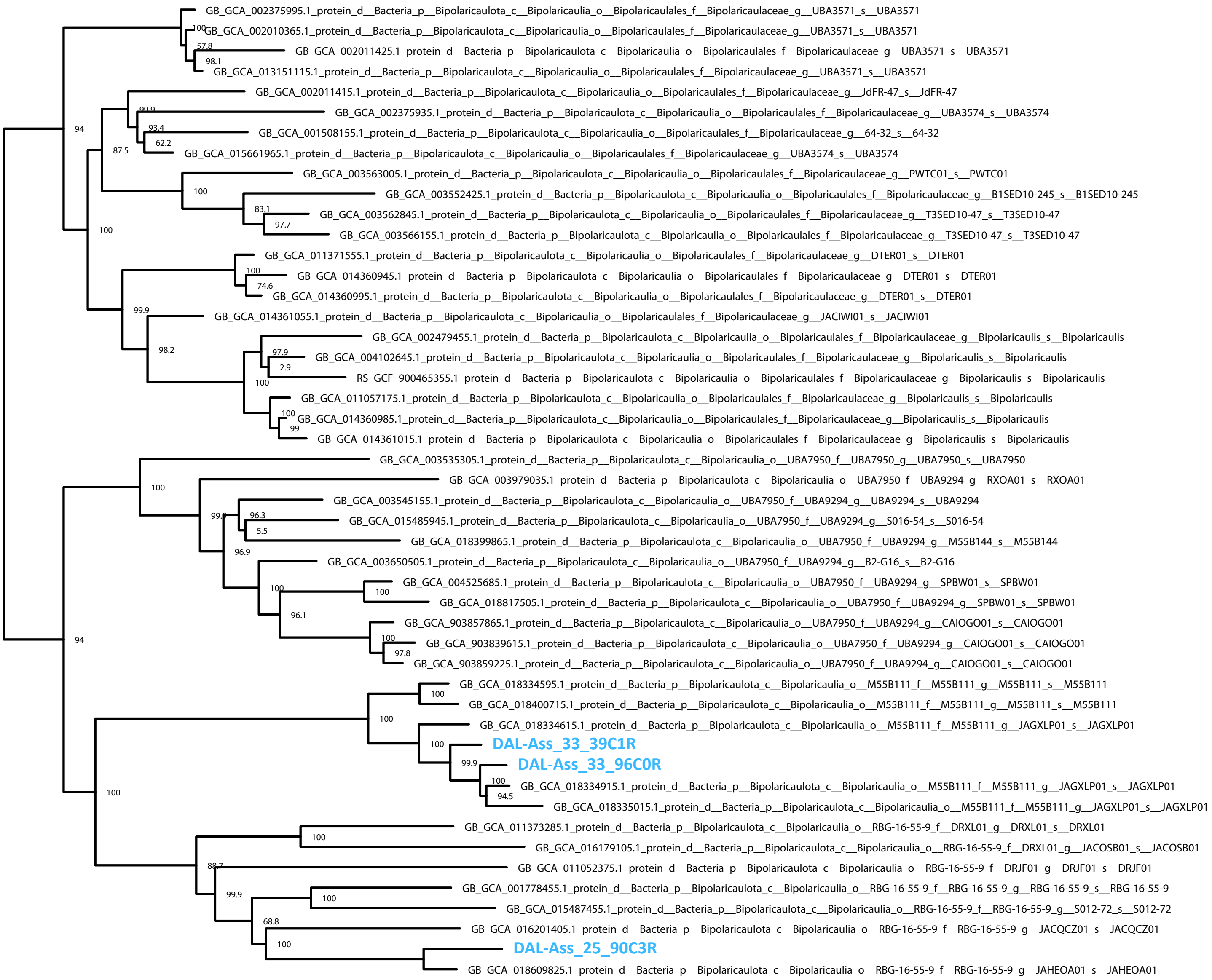


Supplementary Fig.11. Phylogenetic tree of the Bipolaricautota MAGs
